# Components of an ESCRT-independent nuclear envelope assembly pathway

**DOI:** 10.64898/2026.02.01.703137

**Authors:** Emma M. Sydir, M Humam Farra, Abigail L. Whitford, Shea Hinojosa, Pei-Yi Kao, Joao A. Paulo, Sharan Swarup, C. Patrick Lusk, J. Wade Harper, I-Ju Lee, David Pellman

## Abstract

In many cells, the nuclear envelope (NE) must be reassembled after mitosis and holes in the nuclear membrane must be “sealed.” During NE assembly, the NE-specific adaptor, Cmp7, recruits/activates ESCRT-III proteins to mediate NE sealing. However, recent evidence suggests the presence of additional mechanisms. In a screen using the fission yeast, *S. japonicus*, we recently implicated the ESCRT adaptor, Alx1, and a conserved, but little studied protein, Vid27, in Cmp7-independent NE assembly. Here, we provide direct evidence that Alx1 functions in a Cmp7- and ESCRT-independent NE assembly pathway via positive regulation of Vid27. Consistent with a role in membrane remodeling, Vid27 localizes to sites of postmitotic NE sealing and is essential in *S. japonicus*. Alx1 and Vid27 interact and mutations disrupting the predicted interaction interface abolish Alx1’s enhancement of Vid27 function at the NE. These findings define components of a new Cmp7- and ESCRT-independent NE assembly pathway, advancing our understanding of mechanisms that maintain the integrity of the nucleus.

## INTRODUCTION

The nuclear envelope (NE) is a double lipid bilayer that separates the nucleoplasm from the cytoplasm. This compartmentalization is essential for many aspects of cell function and loss of NE integrity is associated with aging, senescence, and cancer (Bakhoum et al., 2018; Papathanasiou et al., 2023; Spektor et al., 2017; Umbreit and Pellman, 2017; Ungricht and Kutay, 2017). The NE is also highly dynamic and undergoes regular remodeling and reorganization, most strikingly during and after mitosis. Strategies for mitotic NE remodeling span a spectrum, from complete breakdown and reassembly of the NE (“open” mitosis) to segregation of chromosomes within a fully intact NE (“closed” mitosis; Dey and Baum, 2021; Makarova and Oliferenko, 2016; Mori and Oliferenko, 2020; Walsh et al., 2024). Although it is obvious that “open” forms of mitosis involve extensive NE disassembly and reassembly, many examples of “closed” mitosis also involve localized NE assembly at sites where nuclear pore complexes (NPC) have been disassembled (Dey et al., 2020) and/or at sites where the spindle pole body (SPB) was inserted or extruded (Cavanaugh and Jaspersen, 2017; Ding et al., 1997). Thus, NE remodeling is a prominent feature of cell division in many eukaryotes.

The endosomal sorting complex required for transport (ESCRT) membrane fission machinery plays a key role in NE remodeling (Frost et al., 2012; Olmos et al., 2015; Vietri et al., 2015). ESCRT machinery mediates membrane fission in various cellular contexts through a common general mechanism (Christ et al., 2017; Hurley et al., 2025; McCullough et al., 2018; Schöneberg et al., 2016; Scourfield and Martin-Serrano, 2017; Vietri et al., 2020a). First, site-specific ESCRT adaptors, such as ESCRT-I/ESCRT-II complexes or the Bro1-domain containing protein, Alx1 (ALIX), recruit ESCRT-III proteins to target membranes (Christ et al., 2017; Henne et al., 2011; Schöneberg et al., 2016; Vietri et al., 2020a). ESCRT-III proteins then polymerize into filaments, simultaneously recruiting the ATPase Associated with diverse cellular Activities (AAA ATPase), Vps4 (VPS4). Vps4 remodels ESCRT-III filaments to induce membrane bending and ultimately fission (Chiaruttini et al., 2015; Adell et al., 2017; McCullough et al., 2018; Mierzwa et al., 2017; Pfitzner et al., 2020).

The only known ESCRT adaptor for NE remodeling is the highly conserved protein, Cmp7 (CHMP7/Chm7; Bauer et al., 2015; Gu et al., 2017; Lee et al., 2020; Olmos et al., 2016; Pieper et al., 2020; Vietri et al., 2015; Webster et al., 2016). Cmp7 is recruited to the NE by the inner nuclear membrane protein, Lem2 (LEM2/Heh1; Gu et al., 2017; Webster et al., 2016). Lem2 binds and activates Cmp7 to initiate ESCRT-III polymerization which in turn promotes NE sealing (von Appen et al., 2020; Thaller et al., 2019). ESCRT-III proteins also recruit the microtubule-severing AAA ATPase, Spastin, thus coordinating spindle disassembly with NE assembly (Olmos et al., 2015; Vietri et al., 2015).

Consistent with its crucial role in maintaining nuclear integrity, the Cmp7 pathway is highly regulated. During interphase, nuclear export of Cmp7 minimizes Cmp7 binding to Lem2 to prevent inappropriate ESCRT-III polymerization (Di Bona et al., 2024; Thaller et al., 2019; Vietri et al., 2020b). Additionally, Vps4 has been shown to disassemble Lem2-Cmp7-ESCRT-III complexes to prevent the formation of Lem2/Cmp7 aggregates that disrupt nuclear integrity and are associated with aberrant centromere positioning and DNA damage (Di Bona et al., 2024; Gatta et al., 2021; Kornakov et al., 2026; Lee et al., 2020; Pieper et al., 2020). Only upon NE breakdown is sufficient Cmp7 activated by Lem2 to initiate ESCRT-III polymerization for membrane sealing (Thaller et al., 2019; von Appen et al., 2020). ESCRT-III activity is further restricted to telophase because of the inhibitory effect of CDK1 phosphorylation of Cmp7 during mitosis (Gatta et al., 2021). Studies have also implicated coiled-coil- and C2 domain-containing protein B (CC2D1B) in binding to ESCRT-III components and regulating the timing of their recruitment to the NE, although the mechanism of this regulation remains unclear (Martinelli et al., 2012; Ventimiglia et al., 2018).

Unexpectedly, the Cmp7 pathway is not strictly essential. In fungi such as *S. cerevisiae* or *S. pombe* that undergo “closed” mitoses, the absence of Chm7/Cmp7 does not significantly affect viability or nuclear compartmentalization (Ader et al., 2023; Dey et al., 2020; Gu et al., 2017; Webster et al., 2016). In *S. japonicus,* a fission yeast that undergoes a “semi-open” mitosis, deletion of *lem2, cmp7*, or other ESCRT genes is detrimental to growth and NE integrity, but the defect is still only partial (Lee et al., 2020; Pieper et al., 2020). In human cells that undergo an “open” mitosis, RNAi depletion of CHMP7 or ESCRT-III proteins delayed but did not fully prevent re-establishment of nuclear compartmentalization after mitosis (Olmos et al., 2015; Vietri et al., 2015). Moreover, *CHMP7* and *LEMD2* are important, but not essential for viability in a subset of human cell lines (Arafeh et al., 2025; DepMap, 2025). Together, these findings suggest that there are ESCRT-independent mechanisms for the assembly of the NE.

To identify factors required for Cmp7-independent NE assembly, we previously performed a genetic screen for mutations that suppress the NE assembly defects of *S. japonicus cmp7Δ* cells. One of the strongest *cmp7Δ* bypass suppressors was a dominant gain-of-function (GOF) mutation in the ESCRT adaptor, Alx1 (Lee et al., 2020). Alx1 is a highly conserved Bro1-domain containing protein that nucleates ESCRT-III activity during membrane remodeling events such as cytokinetic abscission and HIV budding McCullough et al., 2018; Scourfield and Martin-Serrano, 2017; Schöneberg et al., 2016; Vietri et al., 2020a). Surprisingly, our genetic evidence suggested that Alx1 might have an ESCRT-III-independent function at the NE (Lee et al., 2020). Previous work has raised the possibility that Alx1 could have an ESCRT-independent function in viral budding (Bardens et al., 2011; Popov et al., 2009; Shen et al., 2024). However, the underlying mechanism and implications for the normal physiological function of Alx1 function remain unclear.

Here we provide direct evidence that Alx1 promotes NE assembly via an ESCRT-independent mechanism. We find Vid27, another protein identified in our *cmp7Δ* bypass suppressor screen, is a new interaction partner for Alx1 that is crucial for this ESCRT-independent function. Mutagenesis based on *in silico* structural modeling and biochemical experiments confirm the Vid27 interaction with Alx1 and suggest that Vid27 functions downstream of Alx1 during NE assembly. These findings provide a foundation for understanding the mechanism of a newly identified ESCRT-independent pathway promoting postmitotic NE assembly.

## RESULTS

### A gain of function mutation in *vid27* bypasses *cmp7Δ* nuclear integrity defects

Among the bypass suppressors of *cmp7Δ* that we previously identified was a G67R mutation in *vid27* (Lee et al., 2020). Vid27 was initially implicated in several protein degradation and membrane trafficking processes (Brown et al., 2000; Regelmann et al., 2003; Winters et al., 2017; Wolters and Amerik, 2017), but its function remained unclear. More recent work has suggested Vid27 could have a role in cell division and/or nuclear integrity maintenance (Hirano et al., 2020; Lee et al., 2021; Matthew et al., 2022). This indicated that the *vid27-G67R* mutation we identified could be a specific suppressor of *cmp7Δ* nuclear integrity defects, in line with Vid27 playing a role in Cmp7-independent NE assembly.

As a first step to test this hypothesis, we verified the suppression of *cmp7Δ* growth defects by *vid27-G67R* using colony size analysis after ascus dissection (Fig. 1 A). To determine if *vid27-G67R* specifically improved the postmitotic NE assembly defect of *cmp7Δ* cells, we assessed the accumulation of a nuclear import reporter (GST-GFP-NLS-GFP, hereafter GFP-NLS; Yam et al., 2011) in postmitotic, binucleated cells. Indeed, *vid27-G67R cmp7Δ* cells had improved postmitotic NE integrity compared to *cmp7Δ* cells, as assessed by nucleocytoplasmic compartmentalization (Fig. 1 B and Fig. S1 A). Therefore, *vid27-G67R* is a bona fide bypass suppressor of *cmp7Δ*. Consistent with *vid27-G67R* being a gain-of-function (GOF) mutation, we found that mildly overexpressing wild-type *vid27* using the *Purg1* promoter (Watt et al., 2008; Fig. S1 B) also partially rescued the growth defect of *cmp7Δ* cells (Fig. S1 C). However, although Vid27-G67R shows similar steady state levels to our wild-type *vid27* overexpression (Fig. S1 B), Vid27-G67R suppresses *cmp7Δ* growth defects more efficiently than *vid27* overexpression (Fig. 1 A and Fig. S1 C). Thus, in addition to being overexpressed, Vid27-G67R is a GOF variant that promotes NE integrity independently of Cmp7.

**Figure 1.**
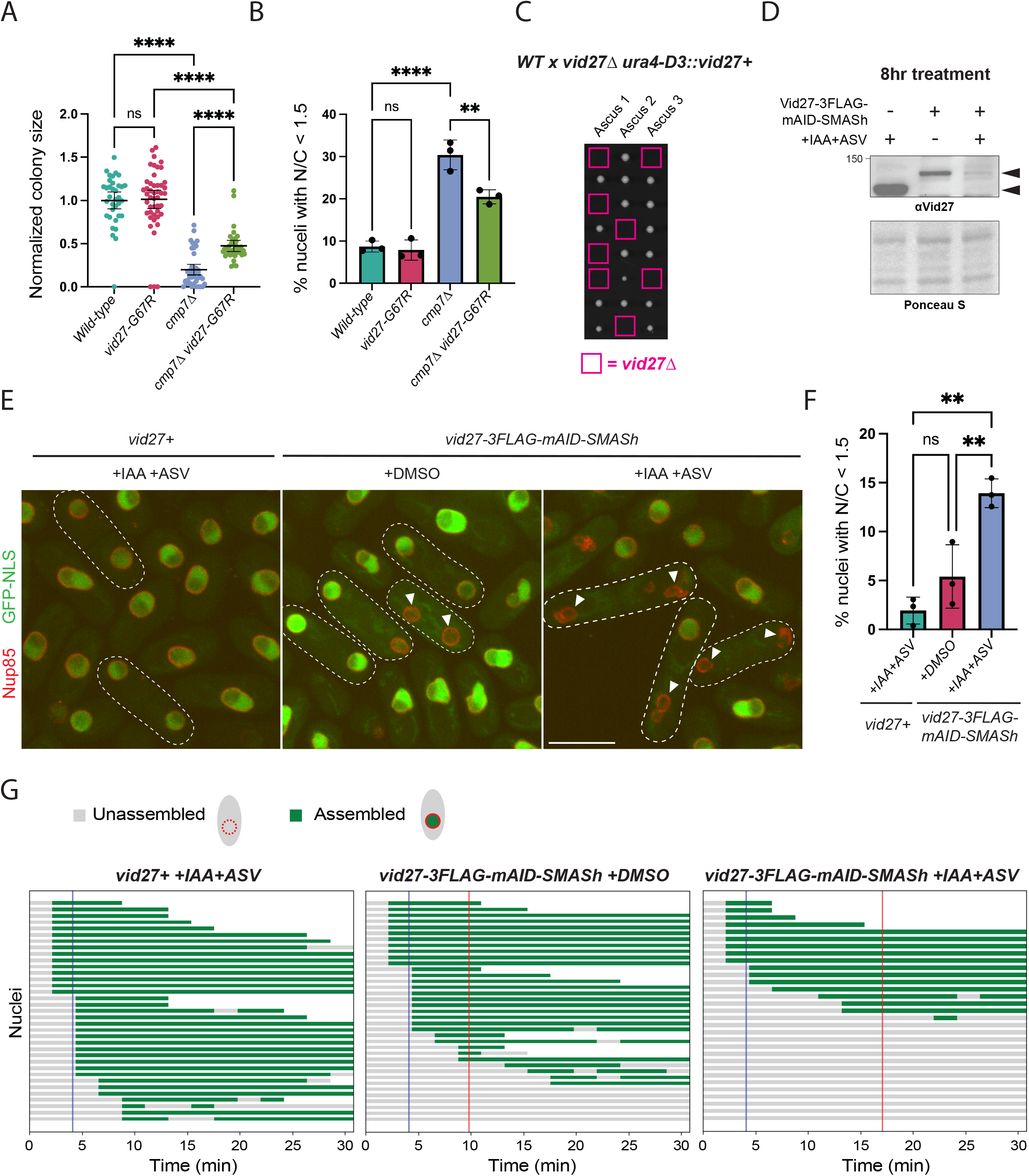
Vid27 is essential in *S. japonicus* and functions in parallel to Cmp7 to promote NE assembly. **(A)** *vid27-G67R* suppresses the growth defect of *cmp7Δ* cells. Normalized colony size after ascus dissection. Means ± 95% CI are shown. ****P≤0.0001; ns, not significant. Brown–Forsythe and Welch ANOVA with Dunnett’s multiple comparison test. **(B)** *vid27-G67R* suppresses the NE integrity defect of *cmp7Δ* cells. Percent of nuclei in binucleated cells with nuclear/cytoplasmic ratio of GFP-NLS<1.5. Means ± 95% CI are shown. n=204-273 cells per strain in each of 3 experiments. **P≤0.01; ****P≤0.0001; ns, not significant; one-way ANOVA and Tukey’s multiple comparison test. **(C)** Vid27 is essential in *S. japonicus*. Representative images of colony growth after ascus dissection of the indicated cross. Note that *S. japonicus* has 8 spores per ascus. Spore positions with the inferred *vid27Δ* genotype in pink squares. **(D-F)** Depletion of Vid27 leads to nuclear integrity defects. **(D)** Western blot showing steady-state levels of Vid27 after 8 h induced degradation (IAA and ASV) and controls. **(E)** Representative images of GFP-NLS and Nup85-mCherry in the indicated cells after 8 h of the indicated treatments. Outlines, binucleated cells; Arrowheads, nuclei with nuclear integrity defects. Scale bar, 10 µm. **(F)** Percent of nuclei in binucleated cells with nuclear/cytoplasmic ratio of GFP-NLS<1.5 after Vid27 depletion. Means ± 95% CI are shown. n=199-498 cells per strain in each of 3 experiments. **P≤0.01; ns, not significant; one-way ANOVA and Tukey’s multiple comparison test. **(G)** Vid27 depletion leads to delays or a block to post-mitotic NE assembly. Traces of a time-lapse series showing the timing of the mitotic loss of NE integrity (time 0) until the end of imaging. Each track represents a daughter nucleus. Green, inferred daughter nuclear reassembly as defined by restoration of NE integrity (GFP-NLS signal at least 50% of the GFP intensity of its mother nucleus from the frame prior to NE breakdown); Gray, time interval where the daughter nucleus does not compartmentalize GFP-NLS; Blue line, average NE assembly time for no degron tag control (4.1 min); Red line, average NE assembly times for degron-tagged strains treated with DMSO (9.8 min) or IAA and ASV (17.1 min). Note that average times are underestimated in Vid27 depletion conditions due to cutoff of imaging time at 30.8 min. n=31-37 nuclei per condition across 3 experiments. Means of distributions compared by permutation test; *vid27+ +IAA+ASV* : *vid27-3FLAG-mAID-SMASh +DMSO,* p=0.0007, *vid27-3FLAG-mAID-SMASh +DMSO* : *vid27-3FLAG-mAID-SMASh +IAA+ASV*, p=0.0068, *vid27+ +IAA+ASV* : *vid27-3FLAG-mAID-SMASh +IAA+ASV,* p<0.0001.

### *S. japonicus* Vid27 is essential and depletion of Vid27 leads to nuclear assembly defects

In other yeasts, such as the highly related fission yeast, *S. pombe*, or the distantly related budding yeast, *S. cerevisiae*, *vid27* is not essential and its loss has little discernable phenotype (Giaever et al., 2002; Hayles et al., 2013; Wysocki et al., 1999). It was therefore notable that we found that *vid27* is essential in *S. japonicus* (Fig. 1 C and Table S1). This raised the possibility that the increased requirement for Vid27 in *S. japonicus* could be due to its “semi-open” mitosis and its consequent increased need for NE remodeling and sealing (Aoki et al., 2011; Heath, 1980; Tanaka and Kanbe, 1986; Yam et al., 2011).

To determine the immediate effects of Vid27 loss, we used a tandem mAID-SMASh degron to deplete Vid27 upon addition of 3-indole acetic acid (IAA) and Asunaprevir (ASV; Lemmens et al., 2018). Even without induced degradation, the Vid27-3FLAG-mAID-SMASh expressing strain showed moderately reduced levels and Vid27 was further depleted, but not completely eliminated, after addition of IAA and ASV for either 4 or 8 hours (Fig. 1 D and Fig. S1 D). Similarly, the Vid27-3FLAG-mAID-SMASh expressing strain exhibited a mild, though not statistically significant, nuclear integrity defect that was significantly exacerbated when IAA and ASV were added (Fig. 1 E and F; and Fig. S1 E-G). Therefore, Vid27 is crucial for maintenance of nucleocytoplasmic compartmentalization.

To directly test if loss of Vid27 delays postmitotic NE assembly, we used time-lapse imaging of Vid27-depleted cells. We defined the NE assembly time as the interval from loss of nuclear-cytoplasmic compartmentalization in mitosis to its reestablishment in interphase daughter nuclei after division. Control cells with wild-type Vid27 treated with IAA and ASV reassembled their nuclei on average within 4.1 min. Even without IAA and ASV-induced degradation, the Vid27-3FLAG-mAID-SMASh expressing strain showed a moderate but significant increase in NE reassembly time to an average of 9.8 min, which was increased to 17.1 min after addition of IAA and ASV (Fig. 1 G and Fig. S2 A-C). Electron microscopy after depletion of Vid27-3FLAG-mAID-SMASh provided additional support for the idea that Vid27 promotes NE membrane sealing following mitosis. Depletion of Vid27 led to ∼4-fold increase in the percent of cells with large (≥150nm) NE gaps (Fig. 2 A and B). Further, the distribution of NE gap length showed a significant shift towards larger gaps in the Vid27-depleted condition (Fig. 2 C). Interestingly, the majority of cells with large NE gaps had only one or two such gaps (Fig. S2 D), consistent with these NE holes being derived from NE assembly failure rather than multisite NE fragmentation. Thus, Vid27 is required for the normal reassembly of intact daughter nuclei following mitosis.

**Figure 2.**
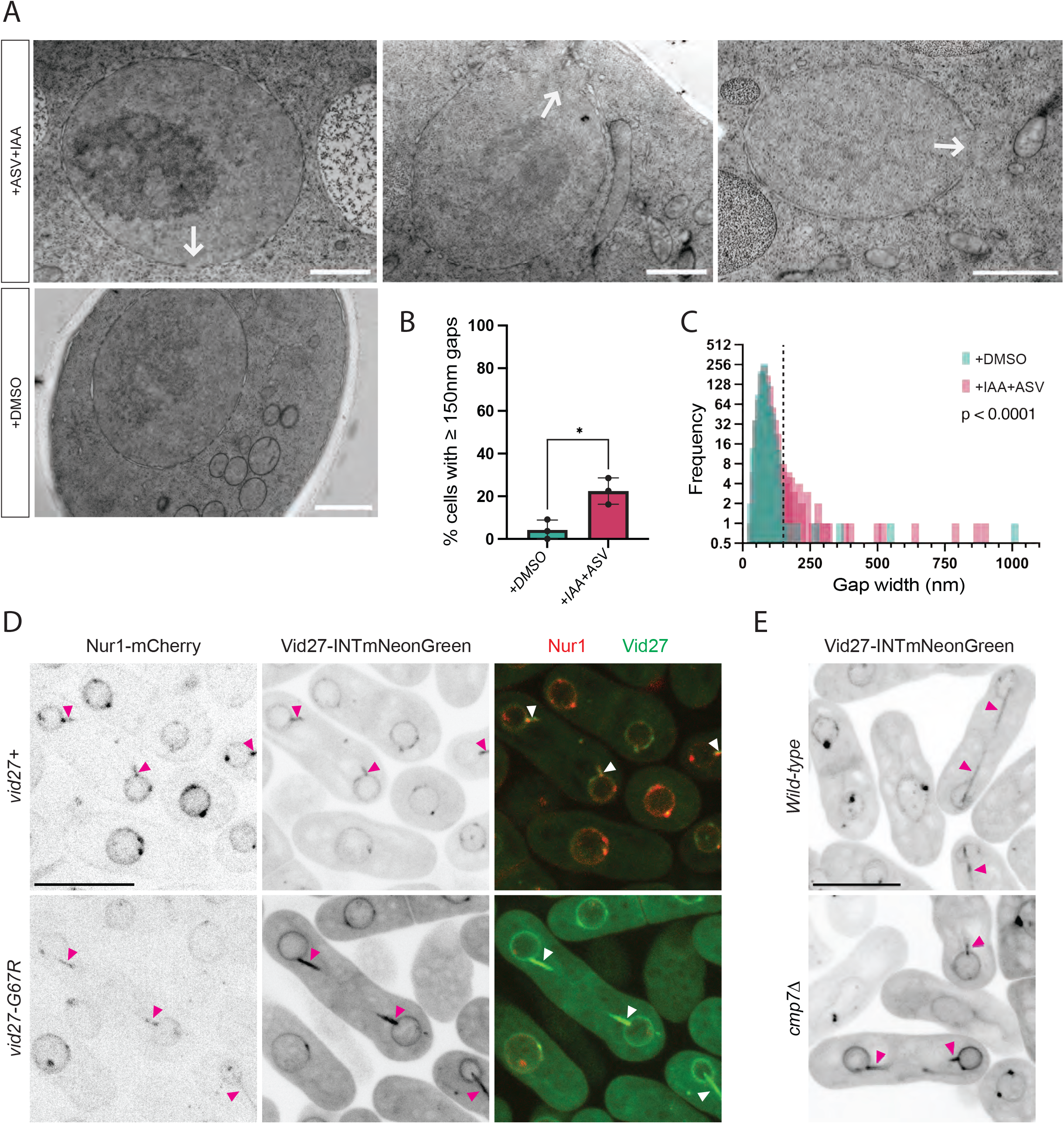
Vid27 localizes to the major sites of NE sealing and is required for proper postmitotic NE assembly. **(A-C)** Depletion of Vid27 results in large NE holes. **(A)** Representative EM images of Vid27-depleted (+ASV+IAA) and control (+DMSO) cells. Arrows indicate ≥150 nm NE gaps. Scale bar, 1 µm. **(B)** Quantification of percent of cells with NE gaps ≥150 nm in Vid27-depleted (+ASV+IAA) vs control (+DMSO) cells. n>90 cells from 3 experiments. Means ± SD are shown. *P≤0.05; Welch’s t test. **(C)** Histogram of NE gap length in Vid27-depleted (+ASV+IAA) vs control (+DMSO) cells. Dashed line indicates 150 nm. n>90 cells from 3 experiments. P<0.001; Kolmogorov-Smirnov test. **(D)** Vid27 and Vid27-G67R localize to sites of postmitotic NE sealing. Shown are representative images of *S. japonicus* cells expressing Vid27-INTmNeonGreen or Vid27-G67R-INTmNG. **(E)** Vid27 localization to “mitotic tails” is independent of Cmp7. Shown are representative images of WT or *cmp7Δ* cells expressing Vid27-INTmNeonGreen. For A & B: Arrowheads, Vid27/Nur1 at mitotic tails; Scale bar, 10 µm.

### Vid27 localizes to the major sites of postmitotic NE remodeling and facilitates membrane sealing in *S. japonicus*

Vid27 could be directly involved in reassembly of the NE or promote postmitotic nuclear compartmentalization indirectly. Consistent with a direct role, Vid27 homologs in other organisms have been reported to be enriched at the NE (Ding et al., 2000; Lee et al., 2021). To assess the localization of the *S. japonicus* protein, we tagged Vid27 at its C-terminus with mNeonGreen (mNG). However, this fusion compromised Vid27 function, as evidenced by *vid27-mNG* being synthetically lethal with *cmp7Δ* (Table S1). Notably, this synthetic lethality implies that Vid27 functions in parallel with Cmp7, consistent with our other data.

We were able to design a more functional Vid27/mNG chimera by introducing the coding sequence for mNG into a segment of *vid27* encoding an unstructured loop (following F429, hereafter Vid27-INTmNG). By contrast with the C-terminal mNG construct, *vid27-INTmNG* did not show synthetic lethality with *cmp7Δ* cells (Table S1).

Using this more functional chimera, we evaluated the localization of Vid27. Vid27-INTmNG exhibited striking accumulation at the sites of postmitotic NE assembly (mitotic “tails”) as well as clear, but less extensive, accumulation at the remainder of the NE (Fig. 2 D). This localization pattern was highly penetrant, with 99% (511/516) of mitotic tails showing an accumulation of Vid27. The localization of Vid27 was not affected by the *vid27-G67R* mutation or by deletion of *cmp7* (Fig. 2 D and E). Thus, Vid27 heavily accumulates at the major site of postmitotic NE sealing, consistent with a direct function in the assembly of the NE.

In addition to localization throughout the NE with enrichment at mitotic tails, Vid27-INTmNG formed NE-associated puncta that did not consistently colocalize with spindle pole bodies (only 2.63% of Vid27 puncta showed overlap with the SPB; Fig. 2 D and E; Fig. S2 E and F).

The lack of Vid27 concentration at SPBs is consistent with other evidence that Vid27 is unlikely to have a major function at the SPB. Vid27 is not essential in *S. pombe* (Hayles et al., 2013) despite the fact that *S. pombe* undergoes SPB insertion/extrusion (Cavanaugh and Jaspersen, 2017; Ding et al., 1997). Moreover, in *S. pombe*, loss of Cmp7 prevents NE sealing at the SPB, although the compartment remains intact, likely because SPB proteins plug the gap (Ader et al., 2023). This inability to seal the SPB hole in the absence of Cmp7 and the ability of SPB proteins to maintain nuclear compartmentalization even in the presence of a gap, suggests that, for SPB sealing, there may not be a parallel sealing mechanism that would require Vid27. Therefore, although the SPB and other sites of NE remodeling have some shared components (Ader et al., 2023), Vid27 appears to be relatively more important for NE assembly at mitotic tails than for SPB insertion or extrusion.

### The essential function of Vid27 is contained within its two N-terminal PH-like domains

From homology and AlphaFold2 modeling, *S. japonicus* Vid27 is predicted to have N-terminal and central PH-like domains (hereafter PH1 and PH2; Fidler et al., 2016; Scheffzek and Welti, 2012) and a C-terminal WD40 repeat-containing β-propeller domain (Fig. 3 A; Jumper et al., 2021; Mirdita et al., 2022; Stirnimann et al., 2010). Each domain is linked to the next by ∼100AA unstructured loops. The *vid27-G67R* mutation falls within PH1 (Fig. 3 A, red). Vid27 is conserved in fungi and plants, and homologs share the general features of one or two PH-like domains linked to a C-terminal β-propeller domain.

**Figure 3.**
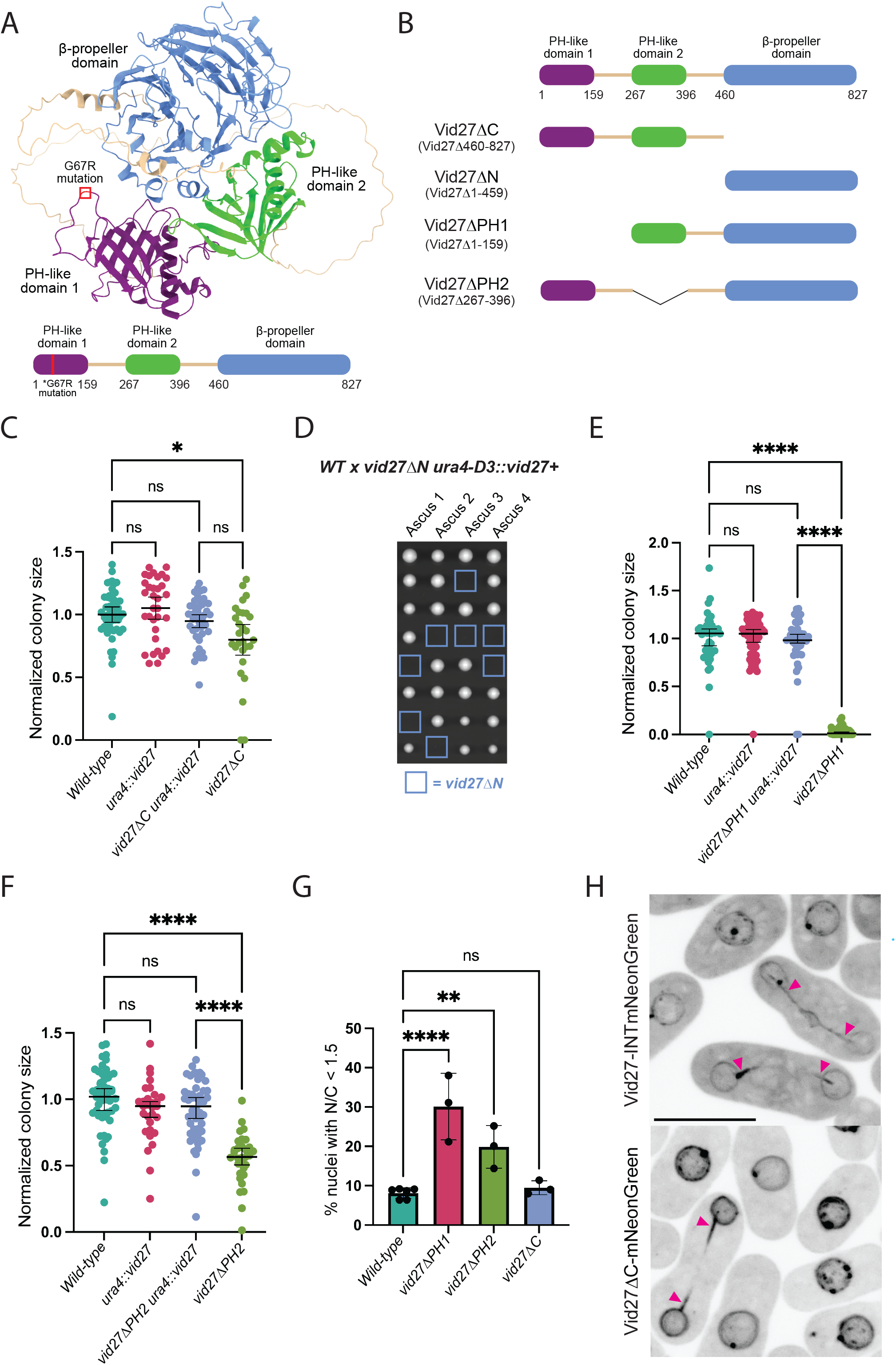
The essential function of Vid27 is contained within its N-terminus, containing its two PH-like domains. **(A)** AlphaFold2 model and cartoon of Vid27 structure. The G67R suppressor mutation is shown in red. **(B)** Design of Vid27 truncations. **(C)** The C-terminus of Vid27 is not essential. Normalized colony size after ascus dissection. Means ± 95% CI are shown. *P≤0.05; ns, not significant; Brown–Forsythe and Welch ANOVA with Dunnett’s multiple comparison test. **(D-G)** The PH-like domains of Vid27 are critical for its NE function. **(D)** Deletion of the N-terminal half of Vid27 causes lethality. Representative images of colony growth after ascus dissection of the indicated cross. Spore positions with inferred *vid27ΔN* genotype in blue squares. **(E and F)** Normalized colony size after ascus dissection for strains expressing *vid27ΔPH1* and *vid27ΔPH2*. Means ± 95% CI are shown. ****P≤0.0001; ns, not significant; Brown–Forsythe and Welch ANOVA with Dunnett’s multiple comparison test. **(G)** Percent of nuclei in binucleated cells with nuclear/cytoplasmic ratio of GFP-NLS<1.5. Means ± 95% CI are shown. n=204-298 cells per strain in each of 3 experiments. **P≤0.01; ****P≤0.0001; ns, not significant; one-way ANOVA and Tukey’s multiple comparison test. **(H)** Vid27ΔC has increased NE localization, including at mitotic tails. Shown are representative images of Vid27-INTmNeonGreen or Vid27ΔC-mNeonGreen localization. Arrowheads, Vid27 at mitotic tails; Scale bar, 10 µm.

To identify the regions of Vid27 that encode its essential function, we generated a series of deletions at the endogenous *vid27* locus in a strain containing a “covering” wild-type copy at the *ura4* locus (Fig. 3 B). Genetic crosses were then used to segregate the *vid27* truncations from the covering copy. Despite being the most conserved feature of Vid27, strains expressing a Vid27 variant lacking the C-terminal β-propeller domain (Vid27ΔC) grew only slightly slower than the control (Fig. 3 C). Vid27ΔC showed substantially higher steady state levels than wild-type Vid27 (Fig. S3 A). By contrast, strains expressing a Vid27 variant lacking the whole N-terminus (Vid27ΔN) were inviable (Fig. 3 D and Table S1), despite Vid27ΔN being expressed at higher levels than wild-type Vid27 (Fig. S3 B). Furthermore, strains expressing variants lacking PH1 or PH2 were significantly compromised in growth, with *vid27ΔPH1* showing a more severe defect than *vid27ΔPH2* (Fig. 3 E and F). Vid27ΔPH2 is expressed at comparable levels to Vid27 whereas Vid27ΔPH1 has a reduced steady state level (Fig. S3 B). The highly impaired growth of *vid27ΔPH1* strains could therefore reflect reduced protein levels, compromised protein function, or some combination thereof.

To assess the importance of each domain for NE assembly, we measured postmitotic GFP-NLS accumulation in cells expressing each truncation. Consistent with our cell growth results, *vid27ΔPH1* showed the highest percentage of cells with postmitotic nuclear integrity defects, followed by *vid27ΔPH2* (Fig. 3 G and Fig. S3 C). By contrast, *vid27ΔC* showed no obvious defect in NE assembly (Fig. 3 G). Further, Vid27ΔC localized to the NE and mitotic tails like Vid27 (Fig. 3 H), albeit with a stronger NE signal, likely due to its elevated steady state levels (Fig. S3 A). Together, these results indicate that the N-terminal half of Vid27 (Vid27ΔC), encompassing its two PH-like domains, performs the protein’s essential function.

However, double mutant *vid27ΔC* and *cmp7Δ* strains exhibited synthetic lethality, demonstrating that Vid27ΔC is not fully functional (Table S1). Because Vid27ΔC shows higher steady state levels than wild-type Vid27 (Fig. S3 A), this truncation may not be a simple loss of function variant, complicating the interpretation of this genetic interaction. Nevertheless, the simplest model is that the C-terminal β-propeller domain has a non-essential regulatory role for Vid27 at the NE that becomes essential when the Cmp7 pathway is lost.

### Vid27 interacts with Bro1-domain containing protein ESCRT-III adapter, Alx1

To identify Vid27 binding partners, we used immunoprecipitation and mass spectrometry (MS) in strains expressing FLAG-tagged Vid27, Vid27-G67R, or Vid27ΔC (Fig. 4 A and B; Fig. S3 D; Table S2). As a validation of the approach, we looked for interaction with the Chaperonin Containing TCP-1 (CCT) which is known to facilitate folding of β-propeller domains (Spiess et al., 2004) and has been reported to robustly bind Vid27 in *S. cerevisiae* (Aloy et al., 2004; Dekker et al., 2008). Consistent with this, we saw that all eight subunits of CCT were enriched in Vid27 immunoprecipitates (Fig. 4 A, cyan dots; Table S2), whereas Vid27ΔC immunoprecipitates exhibited a strong reduction of co-purifying CCT (Fig. S3 D, cyan dots; Table S2). Vid27-G67R immunoprecipitates also showed a trend for enrichment of CCT, however, most subunits fell just below the threshold for statistical significance (∼1.3 on log scale) due to variability between replicates (Fig. 4 B, cyan dots; Table S2). Although the cause of this variability between replicates is not clear, within each replicate of the Vid27-G67R immunoprecipitation, CCT is enriched relative to the untagged control (Table S2). Therefore, our MS experiments capture the primary known Vid27 interactor.

**Figure 4.**
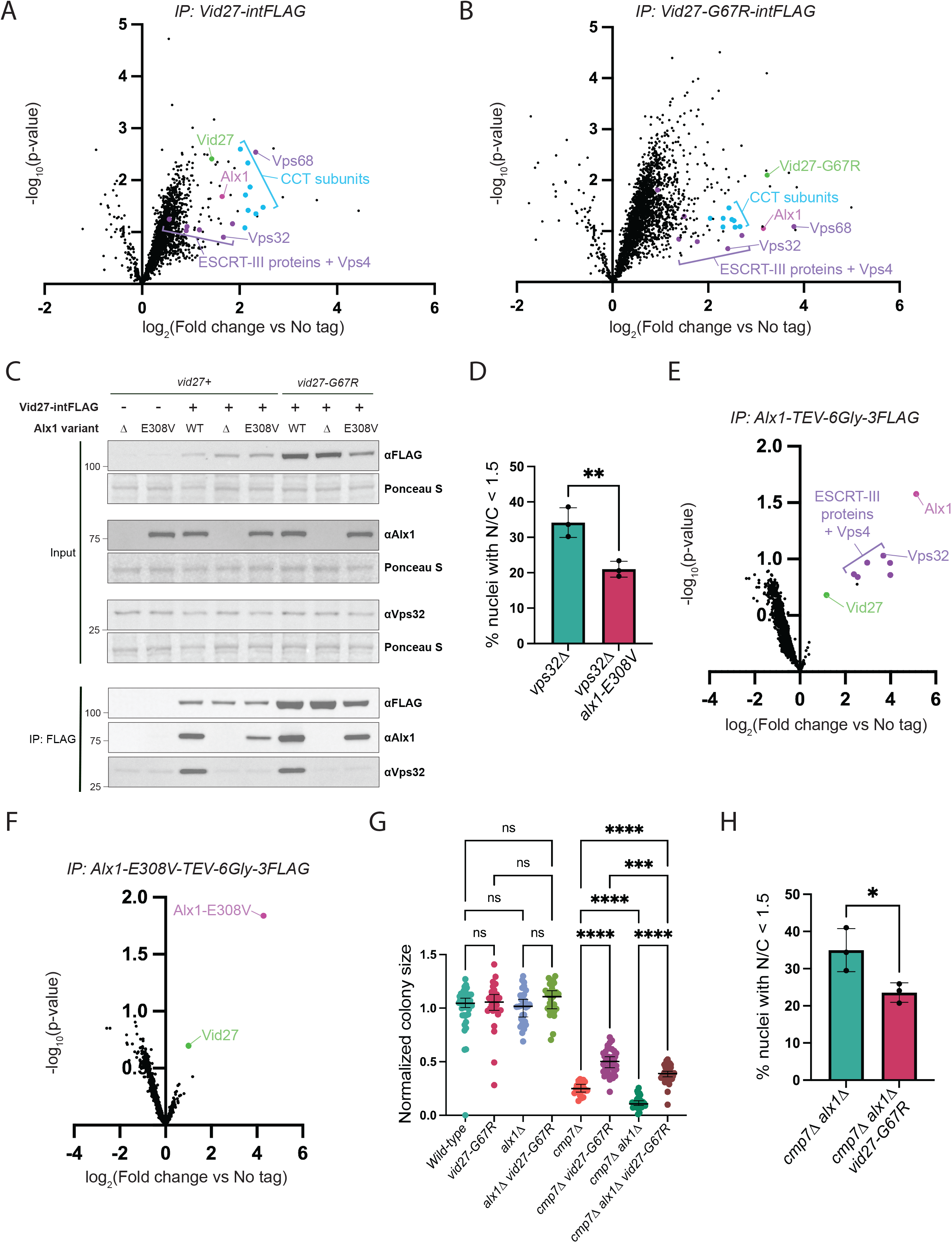
Vid27 and Alx1 interact and promote NE assembly, independent of ESCRT-III. **(A and B)** IP MS indicates that Vid27 and Vid27-G67R interact with Alx1. Volcano plots show TMT-based quantitative MS of Vid27-intFLAG and Vid27-G67R-intFLAG IPs relative to untagged control. Statistical comparisons of 3 replicates by a t-test. **(C)** Vid27 and Vid27-G67R interact with Alx1 and Alx1-E308V, and this interaction bridges Vid27/Vid27-G67R interaction with ESCRT-III. IP of Vid27. IP of Vid27/Vid27-G67R assessed for co-precipitating Alx1/Alx1-E308V and Vps32. **(D)** *alx1-E308V* improves postmitotic NE assembly in cells lacking Vps32. Percent of nuclei in binucleated cells with nuclear/cytoplasmic ratio of GFP-NLS<1.5. Means ± 95% CI are shown. n=200-205 cells per strain in each of 3 experiments. **P≤0.01; one-way ANOVA. **(E and F)** Alx1-E308V appears to interact with Vid27 but lose most interaction with ESCRT-III. Volcano plots show TMT-based quantitative MS of FLAG-tagged Alx1 and Alx1-E308V IPs relative to untagged control. Statistical comparisons of 4 replicates by a t-test. Note that though interactors do not meet the cutoff for statistical significance due to high variability between replicates, signal is enriched for these interactors relative to control in each replicate and key interactors were validated by IP Western blot (Fig. 4 C and Fig. S3 E). Also note that in addition to ESCRT-III, we observe an Alx1 interaction with Sst2, the homolog of an ESCRT-interacting ubiquitin isopeptidase, which is also lost with Alx1-E308V. **(G and H)** *vid27-G67R* suppresses the growth and nuclear integrity defects of *cmp7Δ* strains that lack *alx1*. **(G)** Normalized colony size after ascus dissection. Means ± 95% CI are shown. ****P≤0.0001; ***P≤0.001; ns, not significant; Brown–Forsythe and Welch ANOVA with Dunnett’s multiple comparison test. **(H)** Percent of nuclei in binucleated cells with nuclear/cytoplasmic ratio of GFP-NLS<1.5. Means ± 95% CI are shown. n=212-251 cells per strain in each of 3 experiments. *P≤0.05; one-way ANOVA.

Our MS experiments also pointed to a potential new Vid27 interactor, the ESCRT adaptor, Alx1 (Fig. 4 A and B, magenta dot; Fig. S3 D; Table S2). Interestingly, one of the strongest *cmp7Δ* bypass suppressors from our screen was an allele of *alx1* (*alx1-E308V*, formerly *alx1-m131*; Lee et al., 2020). Alx1 (ALIX in humans) is a Bro1-domain containing protein that promotes ESCRT-III assembly during several membrane remodeling events (e.g., abscission and HIV budding; McCullough et al., 2018; Scourfield and Martin-Serrano, 2017; Schöneberg et al., 2016; Vietri et al., 2020a). Like Vid27, both Alx1 and Alx1-E308V localize to mitotic tails and *alx1Δ* exacerbates the nuclear integrity defect of cmp7*Δ* cells (Lee et al., 2020). Directed co-immunoprecipitation experiments of Vid27, Vid27-G67R, and Vid27ΔC confirmed that all three Vid27 variants consistently and robustly bound Alx1 (Fig. 4 C; Fig. S3 E and F). Thus, two proteins identified in our original bypass suppressor screen localize to mitotic tails and may form a complex, prompting us to investigate the Alx1 suppressor mutation and putative Alx1-Vid27 interaction further.

### *alx1-E308V* establishes the existence of an ESCRT-independent NE sealing pathway

One simple explanation for the bypass of *cmp7Δ* by *alx1-E308V* is that it activates the ESCRT-III pathway downstream of Cmp7. However, several lines of evidence exclude this model. First, Alx1 initiates ESCRT-III activity by recruiting/activating the major ESCRT-III filament-forming protein, Vps32 (Kim et al., 2005; McCullough et al., 2008; Tang et al., 2016; Teis et al., 2008). Yet, our prior work established that, as with *cmp7Δ, alx1-E308V* suppressed the growth defect of *vps32Δ* cells (Lee et al., 2020). Accordingly, we found that *alx1-E308V* also suppressed the NE integrity defect of *vps32Δ* cells (Fig. 4 D and Fig. S4 A), suggesting that *cmp7Δ* suppression by *alx1-E308V* does not require Vps32 recruitment. Second, as expected, immunoprecipitation of wild-type Alx1 followed by MS showed enrichment in Vps32 and other ESCRT-III proteins as well as Vps4 (Fig. 4 E, purple dots; Table S2). Directed co-immunoprecipitation experiments also confirmed the Alx1-Vps32 interaction (Fig. S4 B). By contrast, as assayed by MS, Alx1-E308V almost completely failed to interact with ESCRT-III (Fig. 4 F). In directed co-immunoprecipitation experiments, Alx1-E308V also showed a diminished ability to associate with Vps32, albeit with some variability in the magnitude of this effect between replicates (Fig. S4 B). Consistent with this disruption in binding, the E308V mutation in Alx1-E308V (Lee et al., 2020) is adjacent to the Bro1 domain surface that interacts with Vps32 (McCullough et al., 2008). Therefore, surprisingly, Alx1-E308V’s ability to promote viability and NE assembly is independent of ESCRT-III and the *alx1-E308V* mutation disrupts, rather than enhances, the normal interaction between Alx1 and ESCRT-III. Additionally, Alx1 and Alx1-E308V do not interact with other ESCRT-III “nucleator” proteins, Vps20 and Vps60 (Buysse et al., 2022; Fyfe et al., 2011; Pfitzner et al., 2020, 2023; Fig. 4 E and F; Table S2). Thus, Alx1 regulates a novel ESCRT-independent pathway required for NE sealing.

### Vid27 functions with Alx1 but independently of ESCRT to promote NE sealing

Unlike ESCRT-III proteins, Vid27 was enriched in all replicates of Alx1-E308V immunoprecipitation MS experiments (Fig. 4, E and F, green dot; Table S2) and we were able to validate a robust interaction by Western blot (Fig. 4 C). Therefore, Vid27 could be involved in Alx1’s ESCRT-independent NE assembly pathway. While we also found some enrichment of Vps32 and other ESCRT-III proteins in our Vid27 immunoprecipitation MS experiments (Fig. 4 A and B; Fig. S3 D, purple dots; Table S2), the presence of the Vid27-Alx1 interaction raised the possibility that the Vid27-ESCRT interaction is indirect, bridged by Alx1. Indeed, immunoprecipitation experiments showed that although Vid27 and Vid27-G67R normally co-immunoprecipitate Vps32, in the context of *alx1Δ* or the *alx1-E308V* mutant, the interaction between Vid27/Vid27-G67R and Vps32 is lost (Fig. 4 C). Furthermore, Vid27-G67R was still able to bypass *cmp7Δ* growth and nuclear integrity defects when its binding to Vps32 was disrupted by *alx1Δ* (Fig. 4 G and H; Fig. S4 C).

Our mass spectrometry experiments also indicated that Vid27 could interact with a poorly studied “auxiliary” ESCRT protein, Vps68 (Alsleben and Kölling, 2022; Huh et al., 2003; Schluter et al., 2008; Fig. 4, A and B, Fig. S3 D, purple dot; Table S2). However, colony size analysis showed that Vps68 was not required for either *alx1-E308V* or *vid27-G67R* to suppress growth defects in *cmp7Δ* cells (Fig. S4 D and E). Therefore, the ability of Vid27-G67R to promote NE assembly does not require ESCRT-III or the auxiliary Vps68 protein.

### Vid27 and Alx1 promote NE assembly independent of Lem2 cluster disassembly and independent of Les1

Recent work has implicated several other proteins in Cmp7/ESCRT-independent nuclear integrity maintenance. Lem2 and its binding partner, Nur1, have been shown to form clusters in cells lacking Cmp7/ESCRT-III proteins due to the failure to recruit Vps4 to promote Lem2/Nur1 disassembly (Kornakov et al., 2026; Lee et al., 2020; Pieper et al., 2020). Deletion of either *lem2* or *nur1* abolished cluster formation and improved growth and nuclear integrity in *cmp7Δ* cells, indicating that Lem2/Nur1 clusters are detrimental to nuclear integrity (Lee et al., 2020; Pieper et al., 2020). Disassembly of Lem2/Nur1 clusters is therefore an important mechanism involved in nuclear integrity maintenance.

Interestingly, we observed that in *vid27ΔC* cells, Lem2 formed punctate clusters at the NE (Fig. S5 A), raising the possibility that Vid27 might promote NE assembly primarily by facilitating the disassembly of Lem2/Nur1 clusters. However, several lines of evidence argue against this model. First, although *vid27ΔC* cells robustly formed Lem2 clusters (Fig. S5 A), they exhibited near normal growth and no detectable defect in NE architecture or integrity (Fig. 3 C, G, and H; Fig. S3 C). Second, additional deletion of *lem2* or *nur1* in *vid27ΔC* cells not only failed to rescue the minor growth defect shown by *vid27ΔC* cells, but instead caused lethality (Table S1). Third, *vid27ΔPH1* cells did not form Lem2 clusters (Fig. S5 A) despite exhibiting slow growth and severe defects in nuclear integrity (Fig. 3 E and G; Fig. S3 C). Fourth, Lem2 clusters still form efficiently in *cmp7Δ vid27*-*G67R* double mutant cells, indicating that bypass suppression of *cmp7Δ* by *vid27*-*G67R* does not require cluster disassembly (Fig. S5 B). Finally, in *cmp7Δ vid27*-*G67R* cells, additional deletion of *lem2* decreased rather than increased growth (Fig. S5 C). Therefore, comparing different *vid27* mutants, there is no correlation between NE assembly defects and the formation of Lem2 clusters, indicating that Vid27’s essential function is not connected to the disassembly of Lem2 clusters. These findings are also consistent with our previous work which showed that Alx1 function in promoting nuclear integrity is independent of Lem2 cluster disassembly (Lee et al., 2020).

The NE protein, Les1, was recently shown to limit the sites of localized NE breakdown during *S. pombe* mitosis (Dey et al., 2020). Les1 strongly localizes to mitotic tails, preventing membrane disassembly at these sites to maintain nuclear integrity during cell division (Dey et al., 2020). Because Vid27 shows similar localization to Les1, we considered the possibility that it could function in concert with Les1. However, we found that combining *les1Δ* with *vid27ΔPH2* led to lethality (Table S1), suggesting that Les1 and Vid27 function in parallel pathways to promote nuclear integrity. Additionally, *alx1-E308V* was able to suppress the growth defects of *cmp7Δ* cells even in *les1Δ* strains (Fig. S5 D), indicating that Les1 is not required for Alx1’s NE function. We also tested if *vid27-G67R* could suppress *cmp7Δ* growth defects in the absence of Les1, but found no significant improvement in growth of the *vid27-G67R cmp7Δ les1Δ* strain relative to the *cmp7Δ les1Δ* strain (Fig. S5 E). However, because of genetic linkage between Cmp7, Les1, and Vid27, we were only able to obtain a limited number of triple mutants, which could compromise our sensitivity to detect defects (especially given that *vid27-G67R* is a relatively weak suppressor). These data suggest that Vid27 and Alx1 primarily act in parallel with the Les1 pathway, though we do not exclude the possibility of some shared roles in promoting nuclear integrity. Altogether, Vid27 and Alx1 appear to function independently of known NE assembly and nuclear integrity maintenance mechanisms.

### Vid27 and Alx1 are predicted to form a complex

We used *in silico* modeling with AlphaFold2 to assess if Vid27 and Alx1 could interact directly. Indeed, a predicted interaction was observed between the Proline-rich tail at the C-terminus of Alx1 and Vid27 PH2 domain (Fig. 5 A). This prediction is supported by high AlphaFold2 confidence metrics, with all five models predicting interaction in the same general region and with the top three models showing the same strong interaction interface (Fig. 5 B and C).

**Figure 5.**
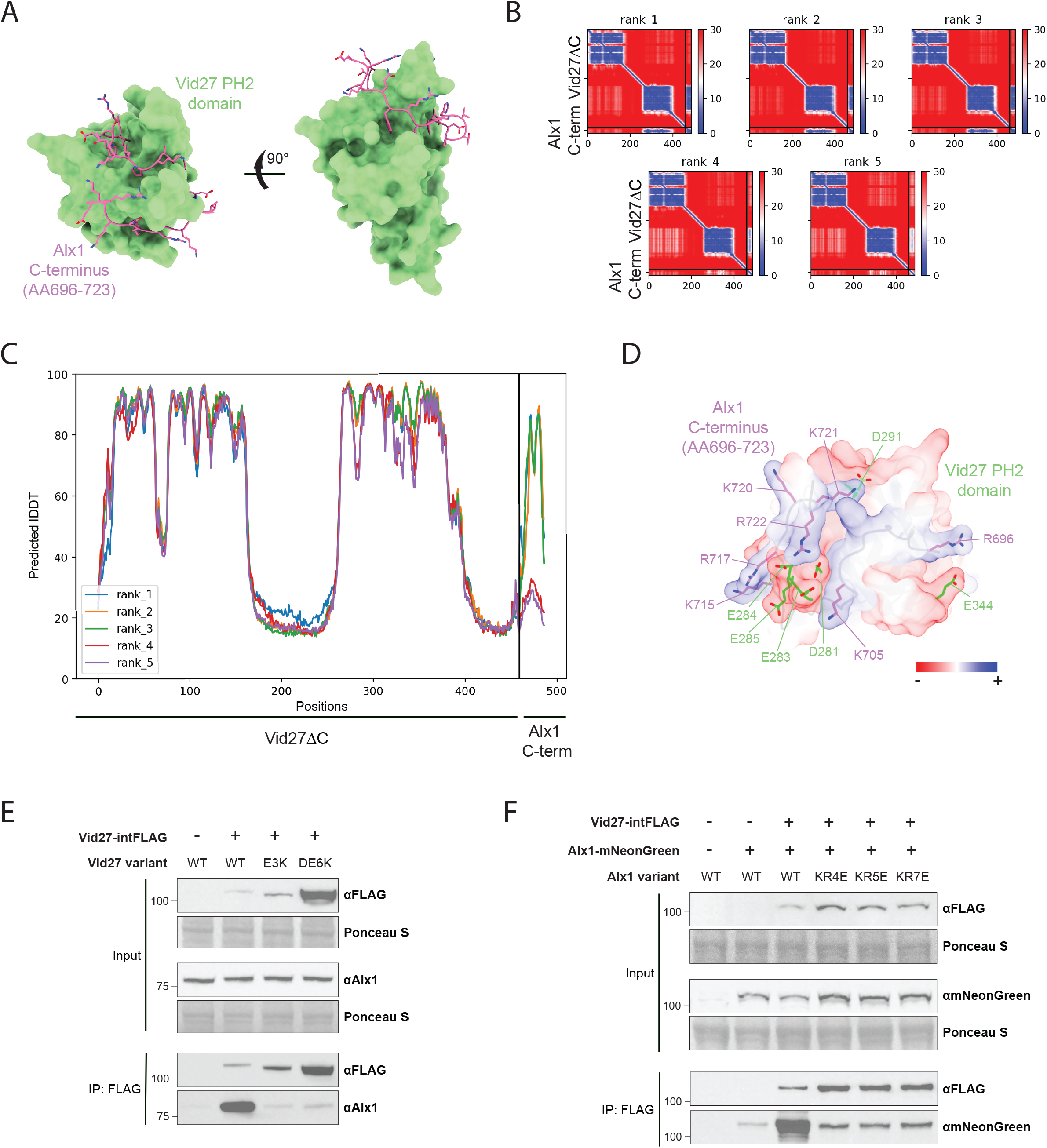
AlphaFold2 prediction and biochemical validation of a Vid27 and Alx1 complex. **(A)** Interaction between the Vid27 PH2 domain (green surface) and C-terminus of Alx1 (magenta sticks) predicted by AlphaFold2. **(B and C)** Confidence metrics for AlphaFold2 predictions of the Alx1-Vid27 interaction. **(B)** Predicted aligned error (PAE) plots. **(C)** Predicted Local Distance Difference Test (pLDDT) plot. **(D)** Electrostatic surface model of Alx1-Vid27 interaction predicted by AlphaFold2. Vid27 acidic residues at the interaction site are labeled in green. Alx1 basic residues at the interaction site are shown in magenta. **(E and F)** Mutations at the predicted Alx1-Vid27 binding interface disrupt the Alx1-Vid27 interaction. **(E)** IP of Vid27 or Vid27 binding surface mutants assessed for co-precipitating Alx1. Vid27-DE6K input band is saturated to allow for visualization of the WT Vid27 input band. **(F)** IP of Vid27 assessed for co-IP of Alx1 or Alx1 binding surface mutants.

This predicted binding interface contains electrostatic interactions, enabling us to design Alx1 and Vid27 variants with charge-swap mutations at the interaction-site residues (Fig. 5 D; Alx1 residues in magenta, Vid27 residues in green). By immunoprecipitation, Vid27 charge-swap mutants, Vid27-E3K (Vid27-E283K,E284K,E285K) and Vid27-DE6K (Vid27-D281K,E283K,E284K,E285K,D291K,E344K), almost completely abolished the interaction with Alx1 despite being present at higher steady-state levels than wild-type Vid27 (Fig. 5 E). Complementary mutations on the Alx1 side of the Alx1-Vid27 interface had a similar effect, as Alx1-KR4E (Alx1-K705E,K715E,R717E,K720E) and Alx1-KR7E (Alx1-R696E,K705E,K715E, R717E,K720E,K721E,R722E) strongly disrupted the Vid27-Alx1 interaction (Fig. 5 F). These findings are consistent with the C-terminal tail of Alx1 binding the Vid27 PH2 domain.

### The ESCRT-III-independent function of Alx1 at the NE requires Vid27

Because Vid27 and Alx1 appear to interact directly, Alx1’s ESCRT-independent function could involve binding to Vid27 in a way that promotes Vid27’s NE assembly function. To test this hypothesis, we determined if the robust bypass of *cmp7Δ* by the Alx1-E308V mutant required Alx1’s ability to interact with Vid27.

We first confirmed by co-immunoprecipitation that Vid27 charge-swap mutants disrupted interaction with Alx1-E308V, similar to how they disrupted the interaction with Alx1 (Fig. 6 A). The requirement for the predicted interaction interface was also supported by immunoprecipitation of Vid27 and assay of its interaction with the complementary Alx1-E308V charge-swap mutants (Fig. 6 B). Thus, Vid27 appears to interact with Alx1-E308V via the same surface predicted to mediate its interaction with Alx1.

**Figure 6.**
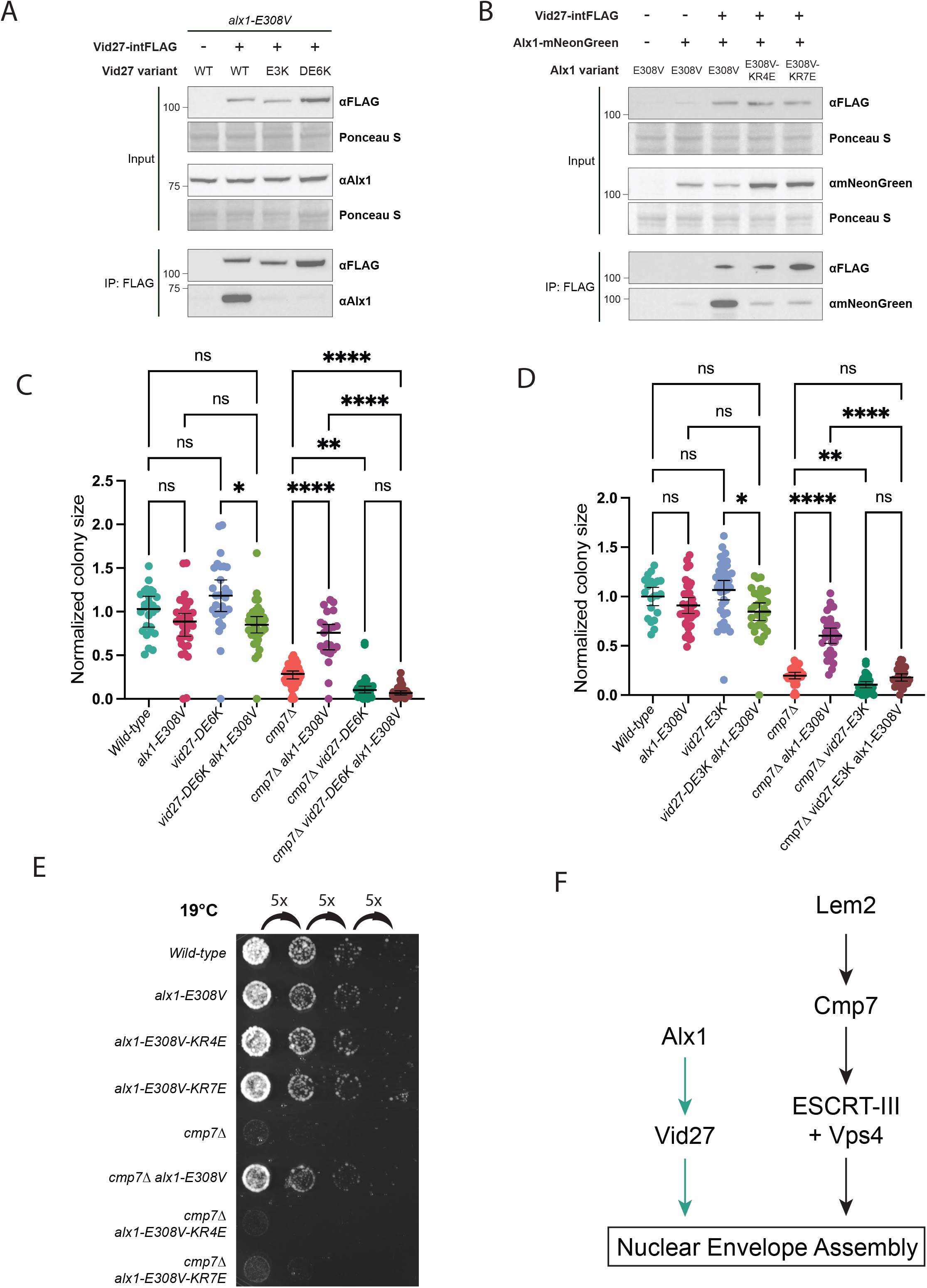
Alx1’s interaction with Vid27 is required for its NE assembly function. **(A and B)** Mutations at the predicted Alx1-Vid27 binding interface disrupt the Alx1-E308V interaction with Vid27. **(A)** IP of Vid27 or Vid27 binding surface mutants assessed for co-IP of Alx1-E308V. **(B)** IP of Vid27 assessed for co-IP of Alx1-E308V or Alx1-E308V binding surface mutants. **(C-E)** *alx1-E308V* cannot suppress the growth and cold sensitivity defects of *cmp7Δ* cells if the Alx1-Vid27 interaction is disrupted**. (C and D)** Effect of Vid27 binding surface mutations on normalized colony size after ascus dissection. Means ± 95% CI are shown. *P≤0.05; **P≤0.01; ****P≤0.0001; ns, not significant; Brown–Forsythe and Welch ANOVA with Dunnett’s multiple comparison test**. (E)** Growth of *alx1-E308V cmp7Δ* at 19°C is abrogated by Alx1 binding surface mutations. **(F)** Proposed model for parallel Alx1-Vid27 and Cmp7/ESCRT mechanisms for NE assembly.

Next, we used colony size analysis to determine if *alx1-E308V* bypass suppression of *cmp7Δ* requires its interaction with Vid27. Indeed, by contrast with the strong suppression of *cmp7Δ* growth defects by *alx1-E308V* (*cmp7Δ alx1-E308V* relative to *cmp7Δ* control), suppression by *alx1-E308V* was abolished by Vid27 charge swap mutants (*alx1-E308V cmp7Δ vid27-DE6K* relative to *alx1-E308V cmp7Δ* control; Fig. 6 C). We saw a similar effect with the *vid27-E3K* mutant (Fig. 6 D). Additionally, *vid27-DE6K cmp7Δ* strains and *vid27-E3K cmp7Δ* strains grew slower than *cmp7Δ* strains (Fig. 6 C and D). This finding mirrors the additive growth defect of *alx1Δ cmp7Δ* strains relative to *cmp7Δ* strains (Lee et al., 2020), again consistent with loss of the Alx1-Vid27 interaction leading to loss of Alx1 function.

We next determined if the Alx1 charge-swap mutations could similarly disrupt *alx1-E308V* suppression of *cmp7Δ* nuclear integrity defects. We previously showed that growing *cmp7Δ* cells at 19°C exacerbates nuclear integrity defects and dramatically slows growth. Furthermore, these phenotypes can be partially rescued by *alx1-E308V* (Lee et al., 2020). Therefore, we determined the extent to which *alx1-E308V-KR4E* and *alx1-E308V-KR7E* could suppress the growth defect of *cmp7Δ* cells at 19°C relative to *alx1-E308V*. In agreement with our model, *alx1-E308V-KR4E cmp7Δ* and *alx1-E308V-KR7E cmp7Δ* strains showed significantly slower growth than the *alx1-E308V cmp7Δ* strain and instead grew comparably to the *cmp7Δ* strain (Fig. 6 E). Altogether, these experiments show that when the Alx1-E308V interaction with Vid27 is disrupted, Alx1-E308V can no longer promote postmitotic NE assembly in the absence of Cmp7. Therefore, the ESCRT-independent function of Alx1 in NE assembly requires its interaction with Vid27.

Because Alx1-E308V requires Vid27 to promote NE assembly, we hypothesized that Alx1 functions upstream of Vid27. This model predicts that Vid27 should be able to promote NE assembly in the absence of Alx1. Indeed, *vid27-G67R* is able to suppress *cmp7Δ* growth and nuclear integrity defects in the context of *alx1Δ*, as *cmp7Δ alx1Δ vid27-G67R* cells showed significantly better growth and postmitotic nuclear integrity than *cmp7Δ alx1Δ* cells (Fig. 4 G and H). Additionally, neither *alx1Δ* nor *alx1-E308V* altered Vid27 localization (Fig. S6 A). Interestingly, *alx1Δ* still partially compromised Vid27 function as evidenced by the *cmp7Δ alx1Δ vid27-G67R* strain growing more slowly than the *cmp7Δ vid27-G67R* strain (Fig. 4 G). Therefore, Vid27’s ability to promote NE sealing is partially but not fully dependent on Alx1, consistent with Alx1 being a non-essential positive regulator of Vid27. Together, our data supports a model in which Alx1 acts as an upstream activator of Vid27 in a newly identified ESCRT-independent NE assembly pathway (Fig. 6 F).

## DISCUSSION

Here, we used a combination of genetics, cell biology, and biochemistry to identify an ESCRT-independent pathway involving the conserved proteins, Vid27 and Alx1. Our work establishes Vid27 as an essential NE assembly factor in this pathway and we further provide evidence that Alx1 is a nonessential positive regulator of Vid27. Our data suggests that this regulation occurs via direct interaction between Vid27 and Alx1, although reconstitution with pure proteins will be important for future characterization of the complex.

By identifying a novel predicted protein complex involved in the process, this study opens up new directions of study in ESCRT-independent NE assembly. We envision two broad classes of mechanism for this ESCRT-independent pathway. First, Alx1 and Vid27 could be components of a yet-to-be identified active membrane fission system. By analogy to other membrane fusion/fission systems, such a system would require a source of energy. In this light, it is interesting to note prior work that implicated SNAREs (Baur et al., 2007; Wang et al., 2013) in NE assembly in *Xenopus* egg extracts. However, although our screens may not be saturated, we did not identify SNAREs in our genetic screen (Lee et al., 2020) or find consistent evidence of SNAREs interacting with Alx1 and Vid27 in our proteomic analysis (Table S2).

Alternatively, rather than being components of an active, energy-requiring fission machinery, Alx1 and Vid27 could promote fission indirectly by bringing membranes into proximity to favor spontaneous NE membrane fission (Ader et al., 2023; von Appen et al., 2020; King et al., 2025; Halfmann et al., 2019, 2023). Given Vid27’s strong enrichment at mitotic tails, it could function by annealing membranes to the spindle as has been proposed for human LEMD2 (von Appen et al., 2020). The positioning of NE membranes in close proximity could facilitate spontaneous fission once the spindle is disassembled. In parallel, Vid27 could also promote spontaneous fission through direct interaction with the NE via its PH-like domains. Recent molecular dynamics simulations suggest that PH domains can thin membranes and disrupt lipid packing, lowering the energy barrier for membrane fission (Baratam et al., 2021).

Such a tethering mechanism could also involve other inner nuclear membrane (INM) proteins identified in our proteomic analyses. For example, we found that Vid27 interacts with the INM protein, Bqt4 (Fig. S6 B; Table S2), an interaction also recently reported in *S. pombe* (Hirano et al., 2023). The interaction may be direct, because Vid27 contains a well-characterized Bqt4 binding motif (Hu et al., 2019). Furthermore, in *S. pombe*, *bqt4Δ* is synthetically lethal with *lem2Δ* (Tange et al., 2016), and the double mutant undergoes extensive NE rupture (Kinugasa et al., 2019). These findings are consistent with Bqt4 functioning in parallel to the Lem2-Cmp7-ESCRT-III pathway to promote NE integrity. How Bqt4 promotes NE integrity remains unclear, but a recent study suggests that Bqt4 promotes the accumulation of phosphatidic acid at the inner nuclear envelope where it would increase negative membrane curvature, promote membrane expansion, and perhaps facilitate spontaneous membrane fission (Hirano et al., 2024). Other studies also support Bqt4 playing a role in regulating membrane composition, possibly through interaction with lipid biosynthesis enzymes (Hirano et al., 2023; Kinugasa et al., 2019). Therefore, Vid27 could work with Bqt4 to generate a NE membrane composition that promotes NE sealing.

Together, our findings establish the existence of an ESCRT-independent NE assembly mechanism and identify protein components of the pathway. Alx1 homologs are found in fungi, plants, and animals, and recent work has shown that Alx1 localizes to sites of NE remodeling in *S. pombe* (Genereux et al., 2026). Vid27 is conserved in fungi and plants, and homologs have been found to localize to the NE in *S. pombe* (Ding et al., 2000) as well as *A. thaliana* (Lee et al., 2021). Although Vid27 does not have an obvious animal homolog, the protein is composed of highly conserved domains and there could be an unrecognized metazoan functional homolog. Therefore, the pathway is likely to be at least functionally conserved and fully elucidating its mechanism will be critical for understanding the cell and disease pathology caused by defects in nuclear integrity.

## METHODS

### *S. japonicus* strain construction

Transformations of *S. japonicus* cells were done using 2–15µg purified DNA fragments per transformation as described in Aoki et al., 2010. DNA fragments for gene targeting were designed as previously described in Bähler et al., 1998, with the modification of using 400-575-bp homologous sequences flanking the insertion site for homologous recombination. These homologous sequences were either amplified from plasmid/genomic DNA or synthesized as double-stranded DNA gene blocks. These sequences were then linked to sequences encoding promoters, fluorescent proteins, and/or selection markers by fusion PCR. Additional gene copies were introduced to the *ura4* locus. To do so, the ORF and 1-kb flanking regulatory sequences were cloned into the pSJK1 plasmid (Aoki et al., 2010) using Gibson Assembly. Alternatively, gene synthesis was used to generate the sequence with the relevant ORF, 1-kb flanking regulatory sequences, and the *kan* gene plus sequences with homology to the *ura4* locus in standard pMA (Thermo Fischer) or pUCIDT (IDT) plasmids. As deletion of *vps32* severely affects growth, the *vps32* deletion strain was generated as described in Lee et al., 2020. Briefly, *kan-Pvps32-vps32-Tvps32* was first integrated at the *ura4* locus after which *vps32* at the endogenous locus was deleted. The resulting strain was then crossed to a WT strain to obtain *vps32Δ* cells. Similarly, to test if *vid27* is essential, *kan-Pvid27-vid27-Tvid27* was first integrated into the *ura4* locus after which endogenous *vid27* was deleted. The resulting strain was then crossed to a WT strain and spores inferred to be *vid27Δ* were assessed for viability. The *alx1-E308V* strain was generated as described in Lee et al., 2020, where it was named *alx1-m131*, based on it being the 131^st^ mutant isolated. Briefly, the endogenous *alx1* ORF was replaced with *Sp\ura4+*, the *S. pombe ura4+* gene (Aoki et al., 2010). In a second round of transformation, *Sp\ura4+* was then replaced with the *alx1-E308V* sequence and counterselected for with 1.5 g/liter 5-fluoroorotic acid (5-FOA) in yeast extract with five supplements (YE5S) media (Furuya and Niki, 2009). The *vid27-G67R* suppressor mutation and *vid27* truncations were introduced to the endogenous *vid27* locus in the same way, except all steps were done in the presence of an extra copy of *vid27* at the *ura4* locus. The resulting strains were then crossed to a WT strain to generate strains lacking the extra copy of *vid27*.

All transformants were selected with selection media and then verified by PCR at the 5′ and 3′ insertion junctions. Point mutations and truncations were further verified by sequencing. All *S. japonicus* strains used in this study are listed in Table S3.

### *S. japonicus* cell culture and genetic crosses

Cells were grown in YE5S at 30°C unless otherwise noted. To assess cell growth at different temperatures, cell cultures were grown to log-phase and then diluted to the same OD600 (0.2–0.4). Using 96-well plates, cells were then serially diluted by 5-fold. Dilutions were then spotted on YE5S plates and grown for 2-3d at 30°C or 5-6d at 19°C and then imaged in Colorimetric mode by a ChemiDoc Imager (BioRad).

For crosses, cells with opposite mating types were combined on sporulation agar and incubated at 30°C for 12–16h or 25°C for 20-24h. Crosses were then dissected with a yeast dissection microscope (Nikon Instruments, Inc., and Micro Video Instruments, Inc.). Asci were incubated at 32-34°C for 6–8h to allow for spores to separate. Spores were then deposited in different positions on a YE5S plate. Dissected spores were incubated at 30°C for 3–6d. Colonies were then streaked and genotyped by replica-plating, PCR, and/or sequencing. *S. japonicus* asci contain eight spores (Furuya and Niki, 2009). However, a complete ascus is indistinguishable under a yeast dissection microscope from an ascus that has lost one or two spores. Therefore, only asci that had six or more spores after dissection were analyzed.

For colony size analysis after ascus dissection, spores were incubated for 3-4d at 30°C. Colonies from germinated spores were then imaged in Colorimetric mode by a ChemiDoc Imager (BioRad) and unbiased colony size analysis was performed using the “Analyze Particles” function in ImageJ/FIJI. For each cross, 20–45 asci were analyzed. Colony sizes were normalized to the average size of WT colonies on the same plate. Only plates containing a minimum of 6 WT colonies for normalization were analyzed. Because of the prominent growth defect of *vps32Δ* cells, we used *vps32Δ::natMX6 ura4-D3::kan-Pvps32-vps32-Tvps32* in crosses to test the interaction between *vps32Δ* and *vid27-m93*. Similarly, as Lem2 has multiple functions and the effect of *lem2Δ* deletion is not fully understood, we used *lem2Δ::natMX6 ura4-D3::kan-Plem2-lem2-Tlem2* in crosses. Only the sizes of colonies lacking the extra copy of *vps32* or *lem2,* respectively, were plotted.

### Immunoprecipitation and immunoblotting

Cells in log-phase (OD600 ≤ 1.2) were pelleted, washed with ddH_2_O, and snap frozen. For each strain, 1-5 g of cell pellet was pulverized (10-12x 2min grind/1min rest cycles at 10cps) in liquid nitrogen using a Freezer/Mill® Dual Chamber Cryogenic Grinder (SPEX SamplePrep, 6875D). Frozen powder was resuspended in 3mL lysis buffer per 1g powder for 2h at 4°C. For pulldown of Vid27-intFLAG/Vid27-m93-intFLAG/Vid27ΔC-6Gly-3FLAG, the following lysis buffer was used: 50 mM sodium phosphate [94.7% dibasic, 5.3% monobasic] pH 8, 100 mM KCl, 10% glycerol, 1% n-Dodecyl-B-D-maltoside (DDM), 1 mM dithiothreitol (DTT), 1mM phenylmethylsulfonyl fluoride (PMSF), and 2x cOmplete EDTA-free protease inhibitor tablet (Roche). For pulldown of Alx1-6Gly-3FLAG, the following buffer was used: 20mM Tris-HCl pH 7.5, 150mM NaCl, 0.5% NP-40, 1mM PMSF, and cOmplete EDTA-free protease inhibitor tablet (Roche). After centrifugation at 17,000g for 15 min, the lysate was incubated with 10-50µL Anti-FLAG® M2 Magnetic Beads (Sigma) at 4°C for 2-4h. Beads were washed 3x with lysis buffer then 1x with exchange buffer (50mM Tris pH 7.5, 150mM NaCl). Proteins were then eluted from beads by 2x 10min of shaking at RT with 250µg/mL 3XFLAG Peptide (Sigma) in exchange buffer. Laemmli SDS-Sample Buffer (Westnet) was then added to eluates.

Input samples were prepared by TCA precipitation of cleared lysates as described in Knop et. al. 1999. Briefly, lysates were diluted to a volume of 1150µL with ice cold ddH_2_O. 150µL of 55% TCA was added and mixture was incubated for 10min on ice. Protein precipitates were pelleted at 17,000g at 4°C for 15min and then resuspended in HU buffer (8M urea, 5% SDS, 200mM Tris pH 6.8, 1mM EDTA, Bromophenol blue, 100mM DTT) by shaking at 65°C for 15min.

Samples were spun down at 17000g for 10min then loaded for SDS-PAGE and Western blotting. Antibodies were used at the following dilutions: Monoclonal anti-FLAG® M2 antibody (Sigma) – 1:3000, mNeonGreen Polyclonal antibody (Proteintech) – 1:500, in-house generated anti-Alx1(AA709-723) peptide antibody – 1:1000, in-house generated anti-Vid27(AA810-827) peptide antibody – 1:1000, in-house generated anti-Vid27(AA388-403) peptide antibody – 1:500, in-house generated anti-Vps32(AA194-210) peptide antibody – 1:1000, anti-mouse IgG (Sigma-Aldrich) and anti-rabbit IgG (Life Technologies) – 1:20,000. Blots were reacted with SuperSignal West Femto Maximum Sensitivity Substrate (Thermo Fisher) and imaged by a ChemiDoc Imager (BioRad). Lysates were stained with Ponceau S (Boston BioProducts) to assess input protein concentration.

### Mass spectrometry sample preparation

Cells were harvested and immunoprecipitation samples were prepared as for immunoblotting with the following modifications. Experiments were scaled up with 20g of cell pellet per strain and 75 µL of beads per sample. For WT Vid27 and Vid27 mutant pulldowns, cell pellets were solubilized in Vid27 MS lysis buffer (50mM sodium phosphate [94.7% dibasic, 5.3% monobasic] pH 8, 100mM KCl, 10% glycerol, 1% DDM, 20mM NaF, 50mM β-glycerol phosphate, 1mM DTT, 1mM PMSF, and 2x cOmplete EDTA-free protease inhibitor tablet [Roche]) and beads were washed with Vid27 MS wash buffer (50mM sodium phosphate [94.7% dibasic, 5.3% monobasic] pH 8, 100mM KCl, 10% glycerol, 1mM DTT, 1mM PMSF, and 2x cOmplete EDTA-free protease inhibitor tablet [Roche]). After elution samples were TCA precipitated by adding 20% TCA and incubating for 45min on ice. Samples were then centrifuged at 17,000g at 4°C for 30min. Supernatant was carefully removed and 10% TCA was added. Samples were then centrifuged at 17,000g at 4°C for 20min. Supernatant was carefully removed and pelleted precipitate was washed 2x with acetone followed by centrifuging at 17,000g at 4°C for 20min. A final wash with methanol was done in the same way. The precipitated proteins were resuspended in 200mM EPPS pH 8.5, digested first by Lys-C overnight at room temperature and later by trypsin (6 h at 37°C). Both enzymes were used at a 1:100 enzyme-to-protein ratio. The samples were then labeled with tandem mass tag (TMTpro) reagents. Acetonitrile was added to a final volume of 30% prior to adding the TMTpro labeling reagent. Labeling occurred at room temperature for 1h. Hydroxylamine was added at a final concentration of ∼0.3% and incubated for 15min at room temperature. Alx1 and Alx1-E308V samples were subjected to fractionation using the high pH reversed-phase peptide fractionation kit (Thermo Fisher) for a final of six fractions. TMTpro-labeled samples were pooled at a 1:1 ratio across all samples. The pooled samples were vacuum centrifuged to near dryness and subjected to C18 solid-phase extraction (SPE) (Sep-Pak, Waters).

### Liquid chromatography and mass spectrometry data acquisition for whole proteome analysis

Vid27 and Vid27-G67R mass spectrometry data were collected using a Orbitrap Exploris480 mass spectrometer (Thermo Fisher Scientific) coupled nLC-1200 liquid chromatograph. Peptides were separated on a 100μm inner diameter microcapillary column packed with ∼35cm of Accucore C18 resin (2.6μm, 150Å, Thermo Fisher Scientific). For each analysis, we loaded ∼2μg onto the column. Peptides were separated using a 150min gradient of 5 to 29% acetonitrile in 0.125% formic acid with a flow rate of 450nL/min. The scan sequence began with an Orbitrap MS1 spectrum with the following parameters: resolution 60K, scan range 350-1350, automatic gain control (AGC) target “standard”, maximum injection time “auto” and centroid spectrum data type. We use a cycle time of 1s for MS2 analysis which consisted of HCD high-energy collision dissociation with the following parameters: resolution 50K, AGC 200%, maximum injection time 96ms, isolation window 0.7Th, normalized collision energy (NCE) 32%, and centroid spectrum data type. Dynamic exclusion was set to automatic. The sample was analyzed thrice with a similar method differing only in the FAIMS compensation voltages (CV) that were used (-40V/-60V/-80V for one analysis and -30V/-50V/-70V for the other two analyses).

Alx1, Alx1-E308V, and Vid27ΔC mass spectrometry data were collected on an Orbitrap Fusion Lumos instrument (using hrMS2-mode). This mass spectrometer was also coupled to a Proxeon NanoLC-1200 UHPLC attached to 100µm capillary column was packed with 35cm of Accucore 150 resin (2.6μm, 150Å; ThermoFisher Scientific) at a flow rate of ∼450nL/min. The scan sequence began with an MS1 spectrum (Orbitrap analysis, resolution 60,000, 350-1350Th, automatic gain control (AGC) target 100%, maximum injection time “auto”). The hrMS2 stage consisted of fragmentation by higher energy collisional dissociation (HCD, normalized collision energy 36%) and analysis using the Orbitrap (AGC 200%, maximum injection time 86ms, isolation window 0.7Th, resolution 50,000). The sample was analyzed twice with a similar method differing only in the FAIMS compensation voltages (CV) that were used (-40V/-60V/-80V for one analysis and -30V/-50V/-70V for the other analysis).

### Mass spectrometry data analysis

Spectra were converted to mzXML via MSconvert (Chambers et al., 2012). Database searching included all entries from the *S. japonicum* UniProt reference Database (downloaded: February 2023). The database was concatenated with one composed of all protein sequences for that database in the reversed order. Searches were performed using a 50-ppm precursor ion tolerance for total protein level profiling. The product ion tolerance was set to 0.03Da. These wide mass tolerance windows were chosen to maximize sensitivity in conjunction with Comet searches and linear discriminant analysis (Beausoleil et al., 2006; Huttlin et al., 2010). TMTpro labels on lysine residues and peptide N-termini +304.207 Da), as well as carbamidomethylation of cysteine residues (+57.021 Da) were set as static modifications, while oxidation of methionine residues (+15.995 Da) was set as a variable modification. In addition, phosphorylation (+79.966 Da) at serine, threonine, and tyrosine residues were also set as variable modifications for phosphopeptide enrichment. Peptide-spectrum matches (PSMs) were adjusted to a 1% false discovery rate (FDR; Elias and Gygi, 2007, 2010). PSM filtering was performed using a linear discriminant analysis, as described previously (Huttlin et al., 2010) and then assembled further to a final protein-level FDR of 1% 5. Proteins were quantified by summing reporter ion counts across all matching PSMs, also as described previously (McAlister et al., 2012). Reporter ion intensities were adjusted to correct for the isotopic impurities of the different TMTpro reagents according to manufacturer specifications. The signal-to-noise (S/N) measurements of peptides assigned to each protein were summed and these values were normalized so that the sum of the signal for all proteins in each channel was equivalent to account for equal protein loading. Finally, each protein abundance measurement was scaled, such that the summed signal-to-noise for that protein across all channels equals 100, thereby generating a relative abundance (RA) measurement.

### Vid27 depletion experiments

Cells in log-phase were diluted to OD600 = 0.1-0.2 and 10µM Asunaprevir (ASV; Cayman Chemical) and 500µM 3-indole acetic acid (IAA; Sigma) or the same volume of DMSO was added. Cells were then grown for 4h or 8h at 30°C. For the 8h treatment, cells were diluted back to OD600 = 0.1-0.2 after 4h to keep cells in log-phase throughout the experiment. At the end of treatment, cells were either imaged or prepared for electron microscopy (EM).

To assess Vid27 protein levels, 1mL aliquots of cells from each condition were collected before drug addition and after 4h or 8h of treatment. Cells were first pelleted and resuspended in 1mL ice cold ddH_2_O. Cells were then lysed as described in Knop et al., 1999 by adding 150µL NaOH lysis buffer (1.85N NaOH, 7.5% β-mercaptoethanol) and incubating on ice for 15min. Samples were then TCA precipitated and prepared for immunoblotting.

### Microscopy and image analysis

For microscopy, cells in log-phase were collected from liquid culture. Except in Vid27 depletion experiments, cells were then centrifuged at 3000g, washed with Edinburgh Minimal Medium (EMM), and imaged in 35mm glass-bottom dishes (MatTek Corporation). Images were collected using a Plan Apo 100×/1.45 NA lens on either a Nikon Ti2 microscope with a Yokogama CSU-W1 spinning disk head, Andor Zyla 4.2 sCMOS camera, Lumencor LED widefield illumination system, and Coherent lasers. Alternatively, we used a Nikon Ti2-E microscope with a Yokogama CSU-W1 spinning disk head, Kinetix sCMOS camera, and Lumencor CELESTA light engine. Microscope software was NIS-Elements. For imaging, a z-focal plane series was acquired every 0.4µm across 8-10µm unless otherwise noted. Maximum-intensity projections of 11-19 in-focus z-slices from the middle of cells are shown. Image analysis and preparation was done in ImageJ/FIJI.

Single z-slices at the mid-plane of each nucleus were used for measuring the nuclear/cytoplasmic ratio of GFP-NLS (Webster et al., 2016). For each cell, the average GFP-NLS intensity across an area of ∼1.3 µm^2^ was measured in both the nucleus and cytoplasm. Background signal was measured and subtracted. 609–1280 nuclei were measured for each strain and/or condition.

Maximum-intensity projections were used for automated colocalization analysis of Vid27 puncta with Pcp1 (SPB marker) using CellProfiler. Briefly, thresholding was used to generate identify Vid27 and Pcp1 puncta and Vid27 puncta were classified as either colocalized or not colocalized with Pcp1 puncta. 86,416 Vid27 puncta were identified and tested for colocalization with Pcp1. Full pipeline details can be found in Table S4.

For nuclear integrity analysis of Vid27 depletion experiments, cells for microscopy were collected from liquid culture containing drugs/vehicle. Cells were then centrifuged at 3000g and washed with the EMM containing drugs/vehicle. Cells were then resuspended in EMM with drugs/vehicle and pipetted onto a gelatin pad with drugs/vehicle, covered with an 18mm round 1.5 coverslip (Warner Instruments), and sealed with nail polish. Gelatin pads were made by combining 0.25g porcine gelatin (Sigma) with 1mL EMM and incubating at 65°C with occasional mixing until dissolved and free of bubbles. Drugs/vehicle were added to the mixture and briefly mixed. 80-100µL of the mixture was pipetted onto a microscope slide and sandwiched with another microscope slide to form a thin “pad.” Pads were stored at RT in a humidified container until ready for use, up to a few hours.

For time-lapse imaging of NE assembly following Vid27 depletion, 4-chamber glass bottom dishes (Cellvis) were treated with 0.5mg/mL sterile Concanavalin A (Sigma) for 5min (1mL per chamber) then aspirated and briefly washed with 200µL of sterile ddH_2_O. 200µL cells in log-phase growth were collected from liquid culture after 4h treatment with drugs/vehicle and added to each chamber. After incubating for 15-30min, cells were gently aspirated, leaving only those adhered to the coverslip, and 1mL media containing drugs/vehicle was added to each chamber. Cells were then imaged every 2.2s for 1h, with a z-focal plane series acquired every 0.4µm across 3µm. Maximum-intensity projections were analyzed for GFP-NLS signal of nuclei during NE reassembly following mitosis. Specifically, the average GFP-NLS intensity across an area of ∼1.3 µm^2^ in the nucleus was measured in each frame starting from the frame before (time -2.2min) NE breakdown (time 0min) until successful NE reassembly or up to 30.8min. GFP-NLS signal was normalized to the intensity of the parent nucleus in the first frame and daughter nuclei were considered reassembled when they had normalized GFP-NLS intensity>0.5. 31-37 nuclei were measured for each strain and/or condition.

### Electron microscopy

Cells were treated with DMSO or ASV and IAA to induce Vid27 depletion as described above. Cells were then pelleted at ∼800 RCF for five minutes and transferred to the 200-μm recess of an aluminum platelet (Engineering Office M. Wohlwend 241). Samples were high-pressure frozen in using an HPM100 (Leica Microsystems). The samples were freeze substituted in 0.1% uranyl acetate and embedded in Lowicryl HM20 (Polysciences) using automated temperature control. Samples were sectioned with a diamond knife (Diatome) mounted on an ultramicrotome (Leica Artos 3D) to a nominal thickness of 250μm. Sections were then collected on carbon-supported 200-mesh copper grids (Ted Pella 01840). Sections were imaged on an FEI Tecnai G2 Spirit BioTWIN. Nuclear envelope gaps were quantified in FIJI.

### Statistical analysis

GraphPad Prism was used for statistical analysis and generating plots. Brown–Forsythe and Welch ANOVA were used to compare colony size because of our assumption that there would be different standard deviations for the colony size of different strains. This was followed by Dunnett’s multiple comparison test. Comparisons of percentage of nuclei with nuclear integrity defects were made using one-way ANOVA with Tukey’s multiple comparison test where appropriate. Comparisons of NE assembly time distributions from time-lapse imaging were done by permutation test. Mass spectrometry data was compared using a two-tailed t-test. Comparison of percent of cells with large NE gaps was done using a simple t-test. Distributions of NE gap lengths were compared using the Kolmogorov-Smirnov test.

## ACKNOWLEDGEMENTS

We would like to thank S. Oliferenko (The Francis Crick Institute, London, UK) and JapoNet (National Institute of Genetics, Shizuoka, Japan) for reagents and the Yale Center for Cellular and Molecular Imaging (CCMI) for EM support. We thank E. Schmid and J. Walter for advice with AlphaFold and G. Brunette for assistance with statistics and data presentation. We also thank V. Denic, D. Lew, D. Moazed, S. Oliferenko, T. Rapoport, S. Shao, and members of the Pellman laboratory for discussion.

E.M. Sydir was supported by the NSF GRFP and NIH/ NIGMS T32 GM008313. The Lusk laboratory was supported by NIH R01 GM105672. The Harper laboratory was supported by AG011085. The Lee laboratory was supported by the National Science and Technology Council, Taiwan, grant number 111-2311-B-007-014 and 113-2311-B-007-009 and Seeding Project 2.0, National Tsing Hua University, Taiwan. D. Pellman is a Howard Hughes Medical Institute investigator, a member of the HMS/Boston Ludwig Center, and this work was supported by NIH grant R37 GM61345.

J.W. Harper is a co-founder of Caraway Therapeutics (a subsidiary of Merck, Inc.) and is a scientific advisory board member for Lyterian Therapeutics. D. Pellman is a member of the Volastra Therapeutics scientific advisory board.

---

**Figure S1.**
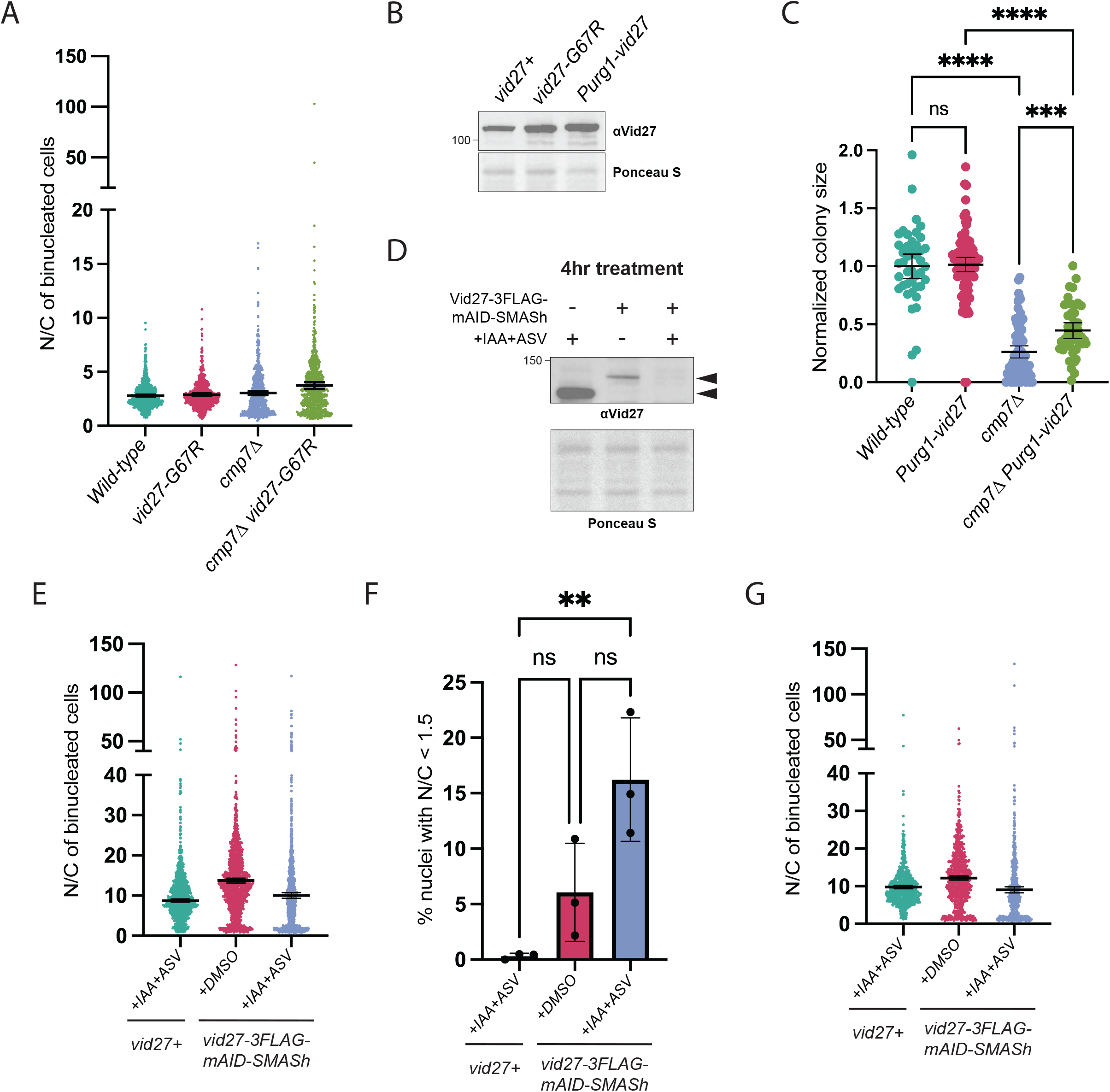
Vid27 is important for nuclear integrity. **(A)** Distribution of nuclear/cytoplasmic ratio of GFP-NLS signal in nuclei of binucleated cells of the indicated genotypes. Means ± 95% CI are shown. **(B)** Western blot showing that Vid27-G67R has moderately increased steady state levels. **(C)** Moderate overexpression of *vid27* suppresses the growth defect of *cmp7Δ* cells. Normalized colony size after ascus dissection. Means ± 95% CI are shown. ****P≤0.0001; ***P≤0.001; ns, not significant; Brown–Forsythe and Welch ANOVA with Dunnett’s multiple comparison test. **(D)** Western blot showing depletion of steady-state levels of Vid27 after 4 h induced degradation (IAA and ASV) relative to controls. **(E)** Scatter plot of nuclear/cytoplasmic ratio of GFP-NLS signal in nuclei of binucleated cells following 8 h of ASV and IAA or DMSO treatment. **(F)** Percent of nuclei in binucleated cells with nuclear/cytoplasmic ratio of GFP-NLS<1.5 after 4 h ASV and IAA treatment to induce Vid27 depletion. Means ± 95% CI are shown. n=202-292 cells per strain in each of 3 experiments. **P≤0.01; ns, not significant; one-way ANOVA and Tukey’s multiple comparison test. **(G)** Distribution of nuclear/cytoplasmic ratio of GFP-NLS signal in nuclei of binucleated cells after 4 h ASV and IAA treatment to induce Vid27 depletion. Means ± 95% CI are shown.

**Figure S2.**
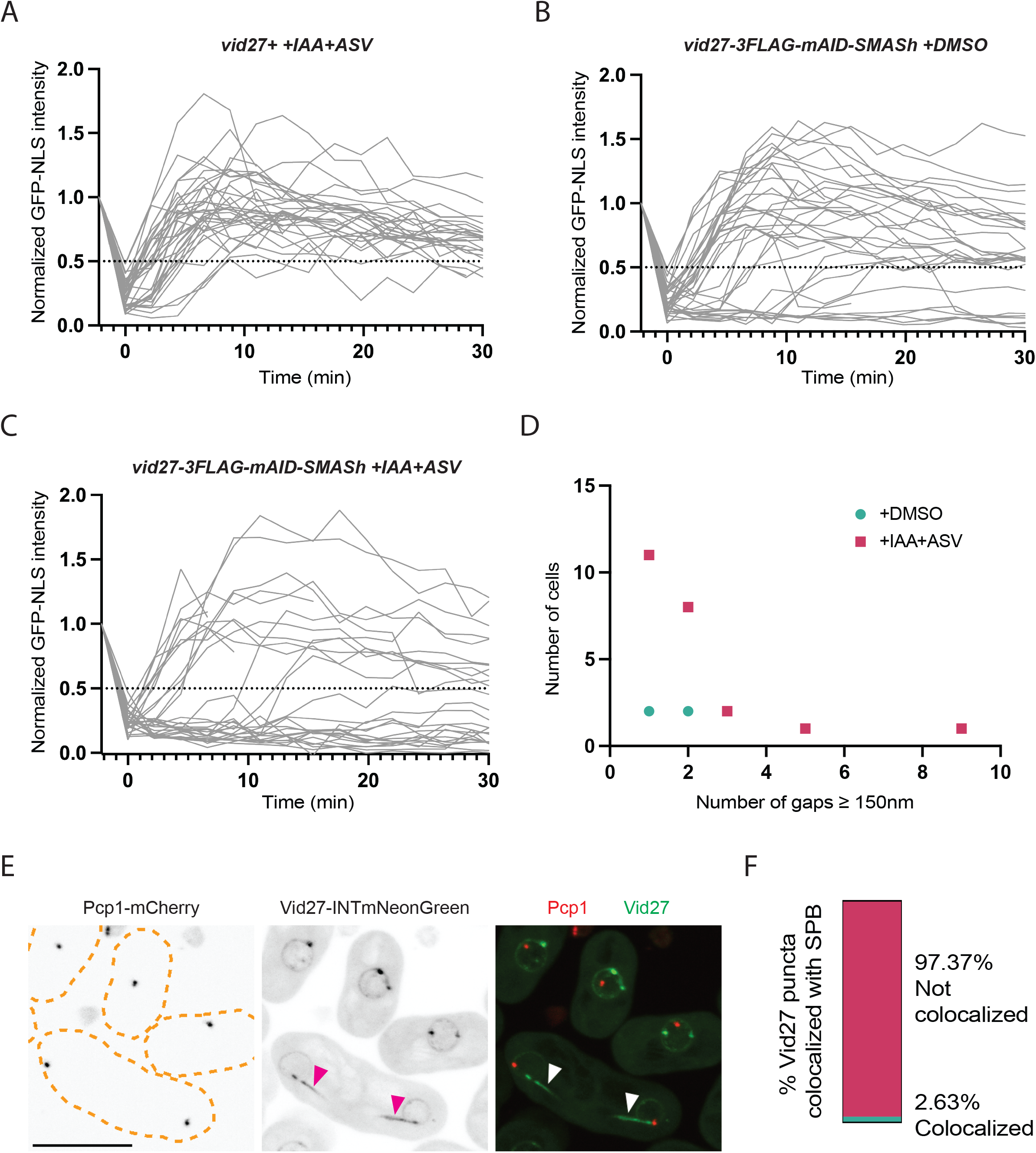
Vid27 primarily functions in NE assembly at mitotic tails. **(A-C)** Time-lapse traces of GFP intensity in each daughter nucleus normalized to GFP intensity in the mother nucleus in the frame before NE breakdown at time 0. Dashed line marks 50% GFP intensity threshold. **(D)** Most cells depleted of Vid27 have only 1-2 large NE gaps (≥ 150nm.), suggesting localized NE disruption, not NE fragmentation. Frequency of cells with different numbers of NE gaps ≥ 150nm for Vid27 depleted (+IAA+ASV) and control (+DMSO) cells (from negative stain EM). **(E and F)** Bright Vid27 puncta do not localize to SPBs. **(E)** Representative images of cells expressing Vid27-INTmNG and Pcp1-mCherry. Arrowheads, Vid27 at mitotic tails; Scale bar, 10 µm. **(F)** Quantification of colocalization of Vid27-INTmNG puncta with Pcp1-mCherry.

**Figure S3.**
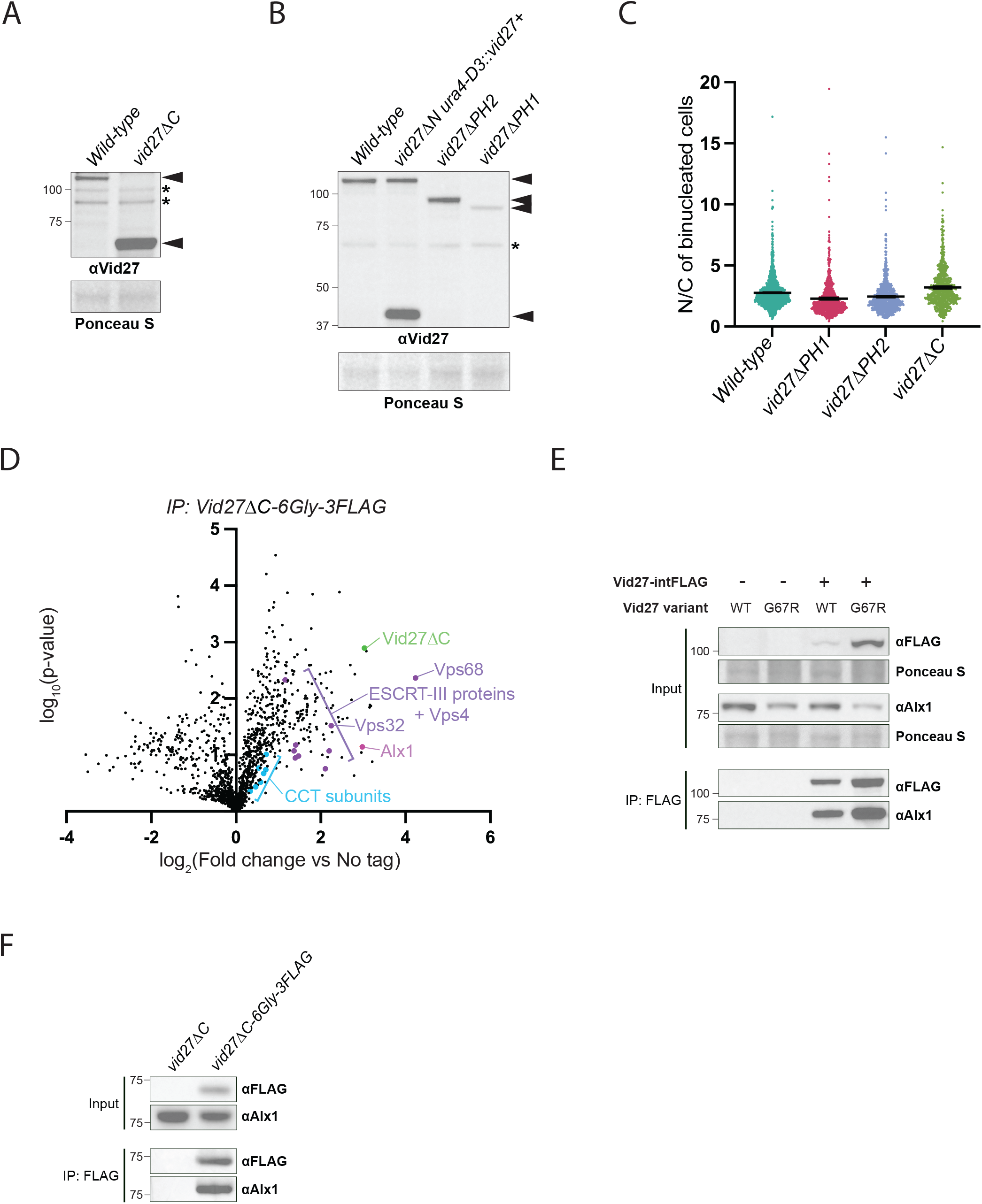
The N-terminal half of Vid27 interacts with Alx1 as part of an ESCRT-independent NE assembly pathway. **(A and B)** Steady-state levels of indicated Vid27 truncations. A covering copy of *VID27* (*ura4-D3::vid27+*) was used to generate a viable strain expressing Vid27ΔN. Note that an antibody to a different Vid27 peptide sequence (AA388-403) was used for Vid27ΔC because this truncation lacks the peptide sequence recognized by our primary Vid27 antibody (AA810-827). Arrowheads, size of Vid27/Vid27 truncations; Asterisks, non-specific bands. **(C)** Scatter plot of nuclear/cytoplasmic GFP-NLS signal in binucleated cells with the indicated genotypes. Means ± 95% CI are shown. **(D)** IP MS suggests that Vid27ΔC interacts with Alx1. Volcano plots show TMT-based quantitative MS of Vid27ΔC-6Gly-3FLAG IPs relative to untagged control. Statistical comparisons of 3 replicates were made using a t-test. **(E)** Western blot of Vid27 and Vid27-G67R co-IP with Alx1. **(F)** Western blot of Vid27ΔC co-IP of Alx1.

**Figure S4.**
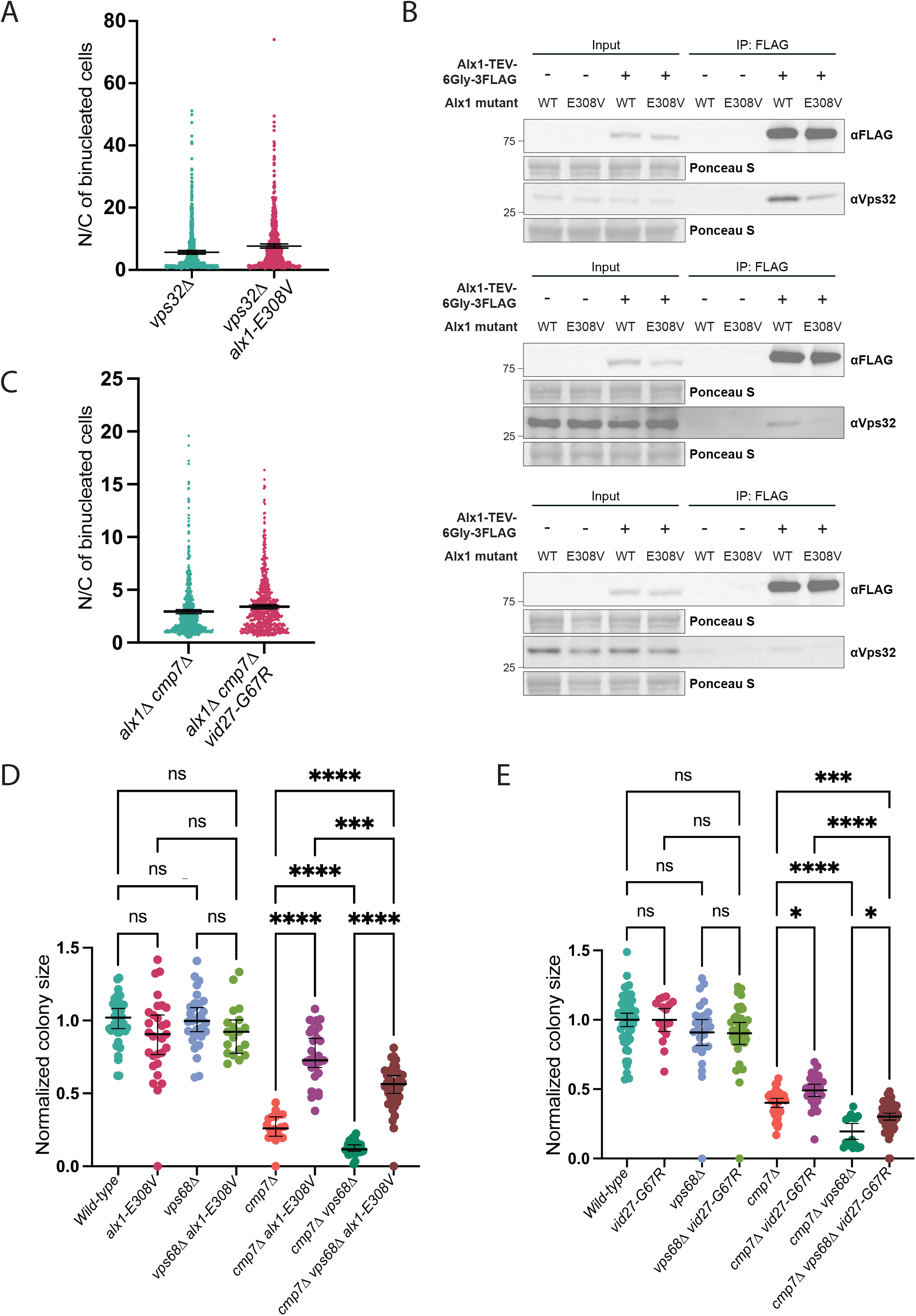
ESCRT-independent NE assembly function of Alx1 and Vid27. **(A)** Distribution of nuclear/cytoplasmic ratios of GFP-NLS intensity in binucleated cells with the indicated genotypes. Means ± 95% CI are shown. **(B)** Reduced interaction between Alx1-E308V and ESCRT-III (Vps32). Shown are three replicate IP/Western experiments **(C)** Scatter plot of nuclear/cytoplasmic ratios of GFP-NLS intensity in binucleated cells with the indicated genotypes. Means ± 95% CI are shown. **(D and E)** *alx1-E308V* and *vid27-G67R* can suppress growth defects of *cmp7Δ* independent of Vps68. Normalized colony size after ascus dissection. Means ± 95% CI are shown. ****P≤0.0001; ***P≤0.001; *P≤0.05; ns, not significant; Brown–Forsythe and Welch ANOVA with Dunnett’s multiple comparison test. Note: Although Vps68 is not required for suppression of *cmp7Δ* by *alx1-E308V* or *vid27-G67R*, there is a negative genetic interaction between *cmp7Δ* and *vps68Δ*, raising the possibility that Vps68 could have some function in parallel to Cmp7.

**Figure S5.**
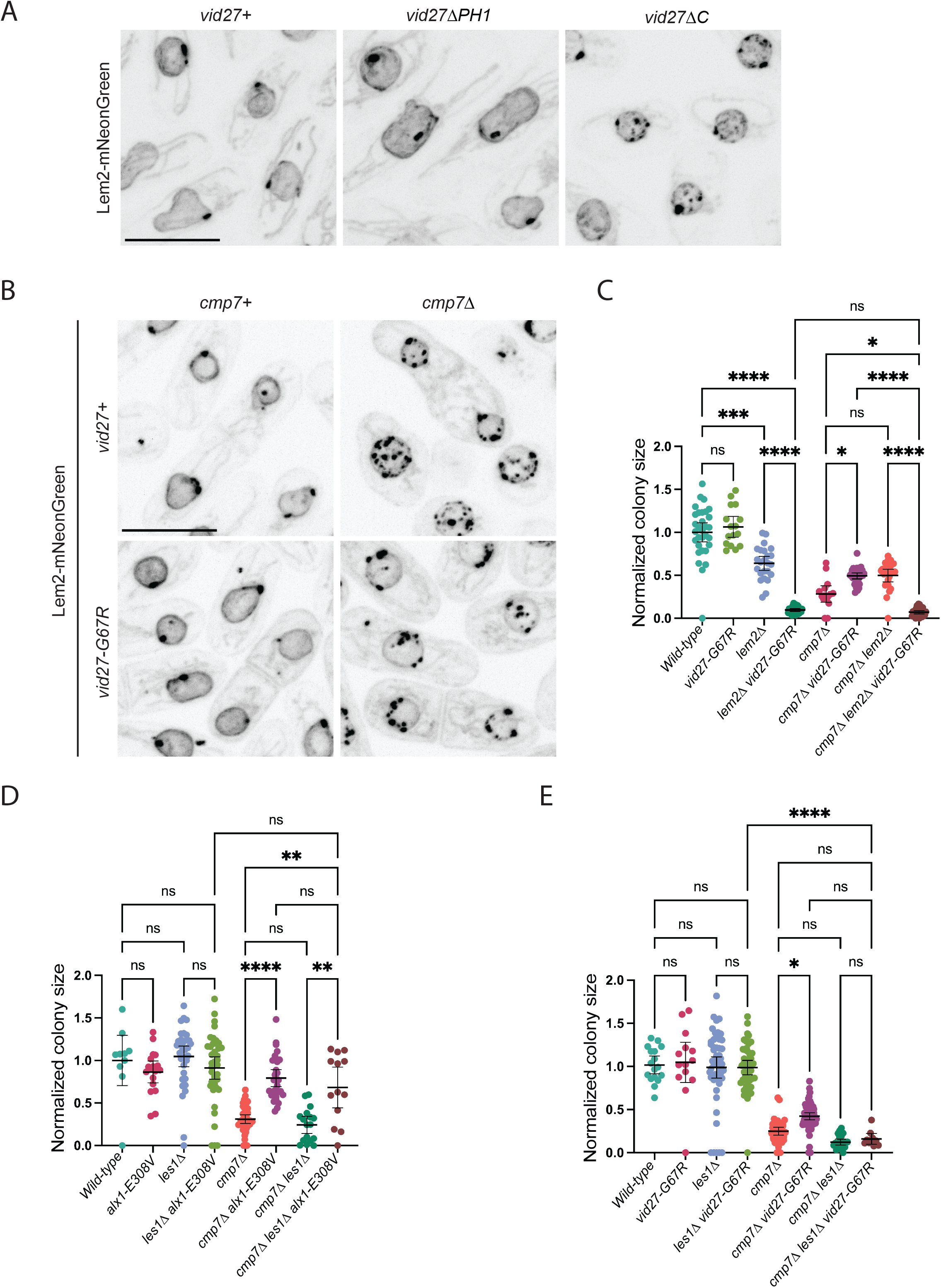
Vid27 promotes NE assembly independent of effects on Lem2 clusters and independent of Les1. **(A)** Lem2 forms clusters in *vid27ΔC* cells but not *vid27ΔPH1* cells. Shown are representative images of cells expressing Lem2-mNeonGreen in the indicated genetic backgrounds. Scale bar, 10 µm. **(B)** *cmp7Δ vid27-G67R* cells have Lem2 clusters. Shown are representative images of cells expressing Lem2-mNeonGreen in *cmp7Δ* and *cmp7Δ vid27-G67R* cells. Scale bar, 10 µm. **(C)** *vid27-G67R* has a negative genetic interaction with *lem2Δ.* Normalized colony size after ascus dissection. Means ± 95% CI are shown. ****P≤0.0001; ***P≤0.001; *P≤0.05; ns, not significant; Brown–Forsythe and Welch ANOVA with Dunnett’s multiple comparison test. **(D)** *alx1-E308V* suppresses *cmp7Δ* growth defects independent of Les1. Normalized colony size after ascus dissection. Means ± 95% CI are shown. ****P≤0.0001; **P≤0.01; ns, not significant; Brown–Forsythe and Welch ANOVA with Dunnett’s multiple comparison test. **(E)** *les1Δ* may compromise the growth of *vid27-G67R cmp7Δ* strains. Normalized colony size after ascus dissection. Means ± 95% CI are shown. ****P≤0.0001; *P≤0.05; ns, not significant; Brown–Forsythe and Welch ANOVA with Dunnett’s multiple comparison test.

**Figure S6.**
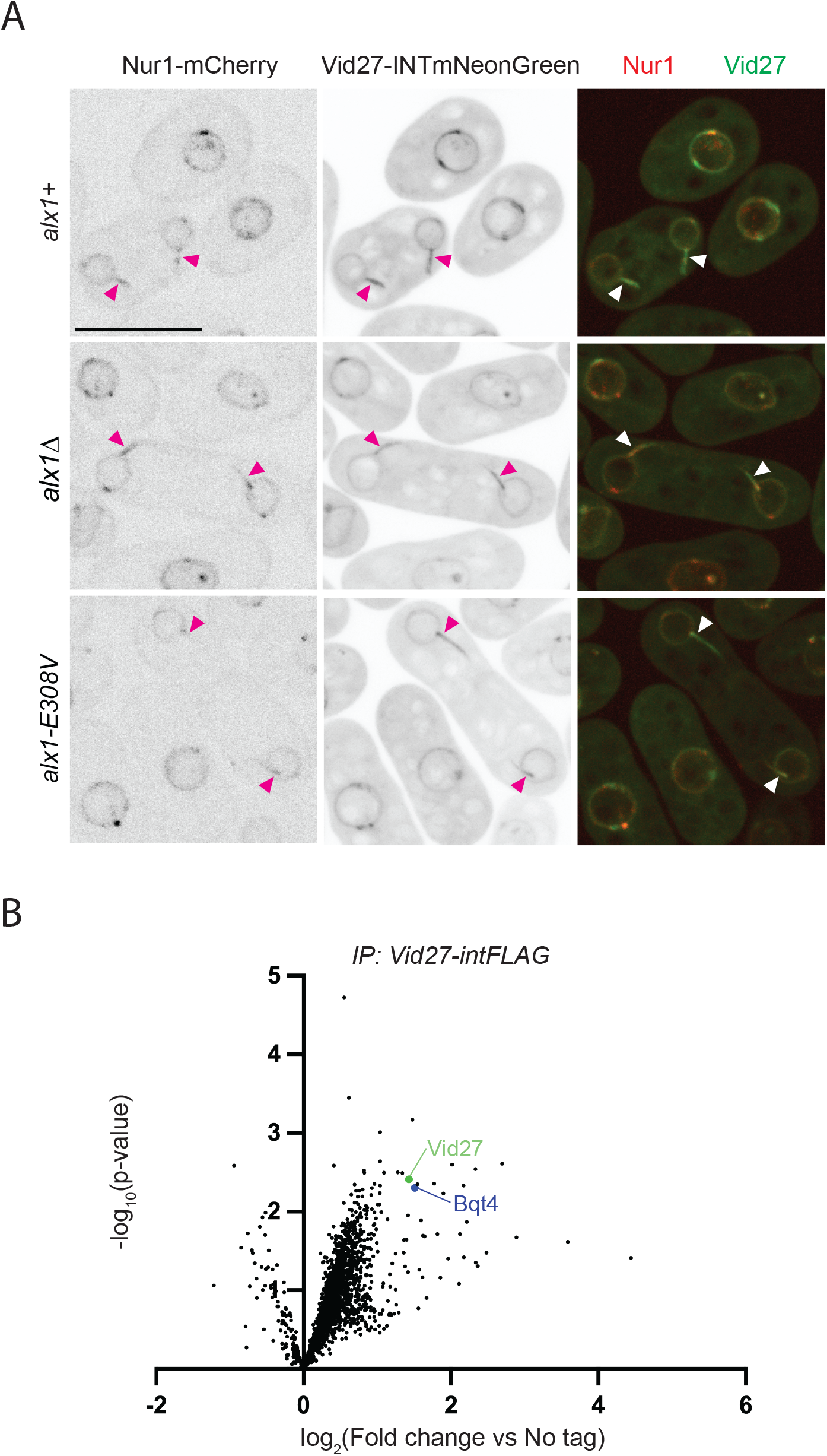
Vid27 localization is not dependent on Alx1, but could be influenced by other interactors. **(A)** Vid27 localization is not affected in *alx1Δ* or *alx1-E308V* strains. Shown are representative images of WT, *alx1Δ,* and *alx1-E308V* cells expressing Vid27-INTmNeonGreen. Scale bar, 10 µm. **(B)** IP MS indicates that Vid27 interacts with Bqt4. Volcano plots show TMT-based quantitative MS of Vid27-intFLAG IPs relative to untagged control. Statistical comparisons of 3 replicates were made using a t test.

